# Breaking the Salinity-Nitrogen Fixation Trade-Off: Engineering A Synthetic Nitrogen-Fixing *Vibrio natriegens* Strain

**DOI:** 10.1101/2025.07.16.665069

**Authors:** Wanjing Wu, Hongzhi Tang

## Abstract

Inefficient nitrogen fixation in legumes under saline stress threatens food security, while conventional symbiotic nitrogen fixation is limited by host specificity and saline stress inhibition. This study engineered *Vibrio natriegens* with nitrogen fixation genes, creating a saline-tolerant, broad-host-range nitrogen-fixing bacteria that enhances plant-microbe interactions through reprogrammed nitrogen metabolism. We demonstrated that engineered nitrogen-fixing *V. natriegen* significantly enhanced soybean growth and nodulation under saline stress. It also activated nitrogenase gene (*nifHDK*) expression in rhizosphere bacteria and increased the abundance of rhizosphere diazotrophs. This study presents a novel approach for developing stress-resilient crop-microbe symbioses, offering a sustainable solution to improve crop growth under saline stress.

## INTRDUCTION

Soil salinization is a major environmental stress that impairs plant growth and reduces crop yield, affecting approximately 400 million hectares worldwide, including 40% of irrigated land.^1^ In legumes, saline stress suppresses rhizosphere diazotroph activity,^2^ impairs rhizobia infection, inhibits nodule formation,^3^ and reduces nitrogenase activity,^4^ collectively diminishing biological nitrogen fixation. Enhancing nitrogen fixation under saline stress is crucial for improving agricultural productivity and addressing food security, yet excessive chemical fertilizer use exacerbates soil salinization and environmental degradation.^5^

In addition to accessing soil nitrogen for growth, legumes can acquire fixed nitrogen through symbiosis with bacteria hosted in root organs known as nodules.^6^ However, conventional symbiotic nitrogen fixation is limited by host specificity and saline stress inhibition. Synthetic biology offers a powerful solution to improve nitrogen fixation under saline stress. Since Dixon’s groundbreaking transfer of the *K. oxytoca* nitrogen fixation (*nif*) gene cluster to *E. coli* in 1972,^7^ precise cross-species transplantation of *nif* gene clusters has been achieved.^8,9,10^ Gene-editing tools now allow the integration of *nif* gene clusters into halotolerant bacterial chassis, creating saline-tolerant nitrogen-fixing engineered bacteria as a sustainable alternative to chemical fertilizers.

As a synthetic biology platform, *Vibrio natriegens* exhibits rapid growth, high halotolerance, and a broad substrate spectrum, making it an ideal chassis for nitrogen fixation.^11,12^ With ribosomal densities reaching 115,000 per cell in the logarithmic phase, *V. natriegens* has significant potential for heterologous protein expression.^13^ Recent advances in genetic tools (e.g., SWAPnDROP,^14^ INTIMATE^15^ and NT-CRISPR,^16,17^ have expanded their engineering capabilities, yet their role in plant growth promotion and nodulation under saline stress remains unexplored.

Here, we engineered *V. natriegens* into a multifunctional plant growth-promoting bacteria (PGPB), which can enhance plant growth and nitrogen assimilation by reprogramming metabolic signaling and activating host-microbiome interactions. Concomitantly, engineered nitrogen-fixing *V. natriegens* (ENF *V. natriegens*) upregulated flavonoids biosynthesis in soybean roots, which modulated the expression of nitrogen metabolism genes and symbiotic nitrogen fixation genes, ultimately boosting soybean nitrogen-fixing capacity under saline stress.

## RESULTS

### Engineering and Functional Characterization of ENF V. natriegens

Native *nif* gene clusters exhibit complex phylogenetic distributions, necessitating the identification of core functional cluster elements. Our engineering strategy prioritized evolutionarily stable *nif* clusters from *Enterobacterales, Bacillales, Pseudomonadales*, and *Rhizobiales*. Redundant genomic regions (e.g., *Pst1307-Pst1312* in *Pseudomonas stutzeri* A1501) were excised,^18^ while essential electron transport components (*rnf/fix* from *Azotobacter vinelandii* DJ) and molybdate transporter genes (*mod*) were incorporated (Fig. 1A).^19^ The *nif* clusters (13-65 kb) from *Paenibacillus polymyxa* WLY78, *Klebsiella oxytoca* M5al, *Azotobacter vinelandii* DJ, and *Pseudomonas stutzeri* A1501 were constructed via *de novo* DNA synthesis or PCR amplification, assembled in yeast, and cloned into host-compatible plasmids. Each gene cluster (13-65 kb) was separately integrated into the chr2_297 site of *V. natriegens* (Vmax) using multiplex genome editing by optimized INTIMATE method, generating ENF *V. natriegens* strains W1, W2, W34, and W78 (Fig. S1). Additionally, a *nif* gene cluster knockout mutant (*Δnif*) was generated in the wild type *V. natriegens* (VCOD-2) as a control (Fig. S2).^15^

**Fig. 1.**
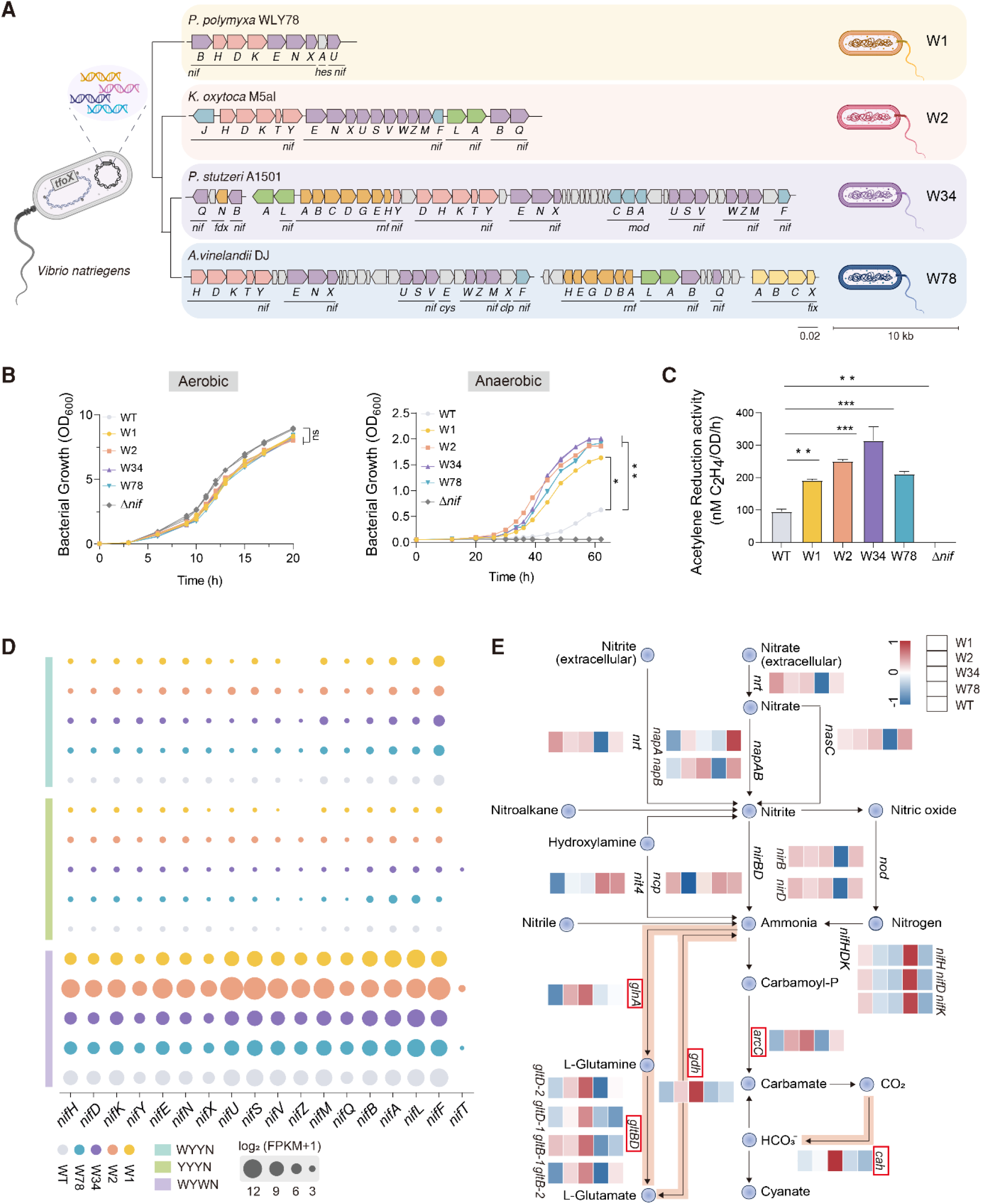
Engineering and functional characterization of ENF *V. natriegens*. **(A)** Alignment of four nitrogen fixation (*nif*) gene clusters from free-living nitrogen-fixing bacteria based on 16S rRNA phylogenetic relationships. Genes are color-coded by function and operon type: red, structural components; purple, cofactor biosynthesis; blue, electron transport; green, regulatory genes; orange, *rnf* and *fix* operons; and grey, unknown nitrogen fixation-related genes. Dots on the DNA line indicate where multiple genomic regions were cloned and combined to form a single plasmid-borne *nif* cluster. The boundaries of all of the *nif* clusters are provided in table. S1, and a complete list of strain genotypes is in table. S2 **(B)** Growth curves of different ENF *V. natriegens* strains under aerobic and anaerobic conditions. Data are presented as the mean ± SD from three biological replicates (\**p* < 0.05, \*\**p* < 0.01, Student’s *t*-test). **(C)** Nitrogenase activity of *V. natriegens* strains carrying native *nif* clusters. Data are presented as mean ± SD from three biological replicates, with error bars indicating standard deviations (\**p* < 0.05, \*\**p* < 0.01, \*\*\**p* < 0.001). **(D)** Effect of different exogenous *nif* gene clusters on endogenous *nif* gene expression in *V. natriegens* under different culture conditions. Bubble size represents gene expression levels; bubble color indicates different strain types; rectangle color denotes culture conditions. **(E)** Expression of genes associated with nitrogen metabolism pathways in different ENF *V. natriegens*. Compared to wild type *V. natriegens*, ENF *V. natriegens* W34 shows upregulated expression of genes associated with the upstream glutamate metabolic pathway. Green highlights indicate relevant pathways and red boxes denote associated genes.

To assess whether these heterologous *nif* clusters were functional in *V. natriegens*, we evaluated growth dynamics and nitrogenase activity across ENF *V. natriegens*. Under aerobic conditions, no discernible differences in growth rates were observed, although the *Δnif* mutant displayed slightly improved growth, suggesting a metabolic burden associated with *nif* cluster maintenance. However, under anaerobic conditions, ENF *V. natriegens* strains exhibited improved growth relative to wild type *V. natriegens* (Fig. 1B). Nitrogenase activity measurements followed a similar trend, confirming that heterologous *nif* clusters enhanced functional nitrogen fixation capability (Fig. 1C).

To investigate the nitrogen metabolic reprogramming induced by *nif* cluster integration, we performed transcriptome analyses under three growth conditions: aerobic with nitrogen supplementation (YYYN), anaerobic without nitrogen (WYWN), and anaerobic with nitrogen supplementation (WYYN). Gene expression profiles displayed marked divergence across these conditions (Fig. S3A). Principal component analysis (PCA) revealed high intra-group consistency and clear inter-group separation (Fig. S3B) (ADONIS: YYYN *R*^*2*^ = 0.91, *p* = 0.001; WYWN *R*^*2*^ = 0.84, *p* = 0.001; WYYN *R*^*2*^ = 0.87, *p =* 0.001), identifying nitrogen availability as a key regulator of native *nif* cluster expression (Fig. S4). Subsequent qPCR analysis confirmed that heterologous *nif* clusters were actively expressed in all ENF *V. natriegens* strains (Fig. S5).

Interestingly, we observed that the integration of *K. oxytoca* M5al *nif* clusters upregulated endogenous *nif* gene expression in *V. natriegens*, while *A. vinelandii* DJ and *P. stutzeri* A1501 *nif* cluster insertions downregulated native *nif* expression (Fig. 1D). This regulatory divergence likely reflects the differing aerobic and anaerobic adaptations of donor organisms, which differentially modulate nitrogen metabolic pathways in *V. natriegens*. To further elucidate how heterologous *nif* clusters influence nitrogen metabolism, we conducted a comparative transcriptomic profiling focusing on nitrogen-associated pathways. Notably, in W2, genes involved in dissimilatory and assimilatory nitrate reduction were downregulated relative to other variants. In contrast, W34 exhibited marked upregulation of glutamate metabolism regulators (*gltBD, gdh, glnA*), carbamate kinase (*arcC*) and carbonic anhydrase (*cah*) gene (Fig. 1E). Meanwhile, expression of amino acid biosynthesis genes was increased (Fig. S6). These findings demonstrate that native *nif* integration enhances both nitrogen fixation capacity and nitrogen metabolic flux in ENF *V. natriegens*.

### Growth promotion of soybean by ENF V. natriegens under saline stress

We observed that the integration of the *nif* gene cluster in ENF *V. natriegens* modulated the expression of genes associated with chemotaxis and cell motility (Fig. S7), which are critical for root colonization capacity in bacteria.^20^ Specifically, *nif* gene cluster integration in W34 upregulated plant growth-promoting genes (Fig. S8) and markedly improved the secretion of phytohormones, including indole-3-acetic acid (IAA) and siderophores (Fig. S9). These findings aligned with soybean seed germination results under saline stress (Fig. S10). Pot experiments further confirmed that inoculation with W34 and W78 significantly promoted soybean seedling growth under saline stress (Fig. S11).

Given that *V. natriegens* lacks nodulation genes, we hypothesized that co-inoculation with *Bradyrhizobium diazoefficiens* USDA110 could enhance root development in legumes.^21^ ENF *V. natriegens* W34/W78 conducted pot experiments were selected (Fig. 2A). Under 500 mM NaCl condition, negative control plants (non-inoculated) displayed severe saline stress phenotypes, including leaf curling and wilting. Soybeans inoculated with USDA110 alone exhibited substantial growth inhibition, with similar suppression observed in the WT+USDA110 group. In contrast, experimental groups co-inoculated with ENF *V. natriegens* (W34/W78) exhibited robust growth (Fig. 2B), with shoot fresh weight increasing by 102.7-284.6%, dry weight elevating by 146-275.7%, and plant height improving by 87-180% relative to the WT+USDA110 group (Fig. 2C-E).

**Fig. 2.**
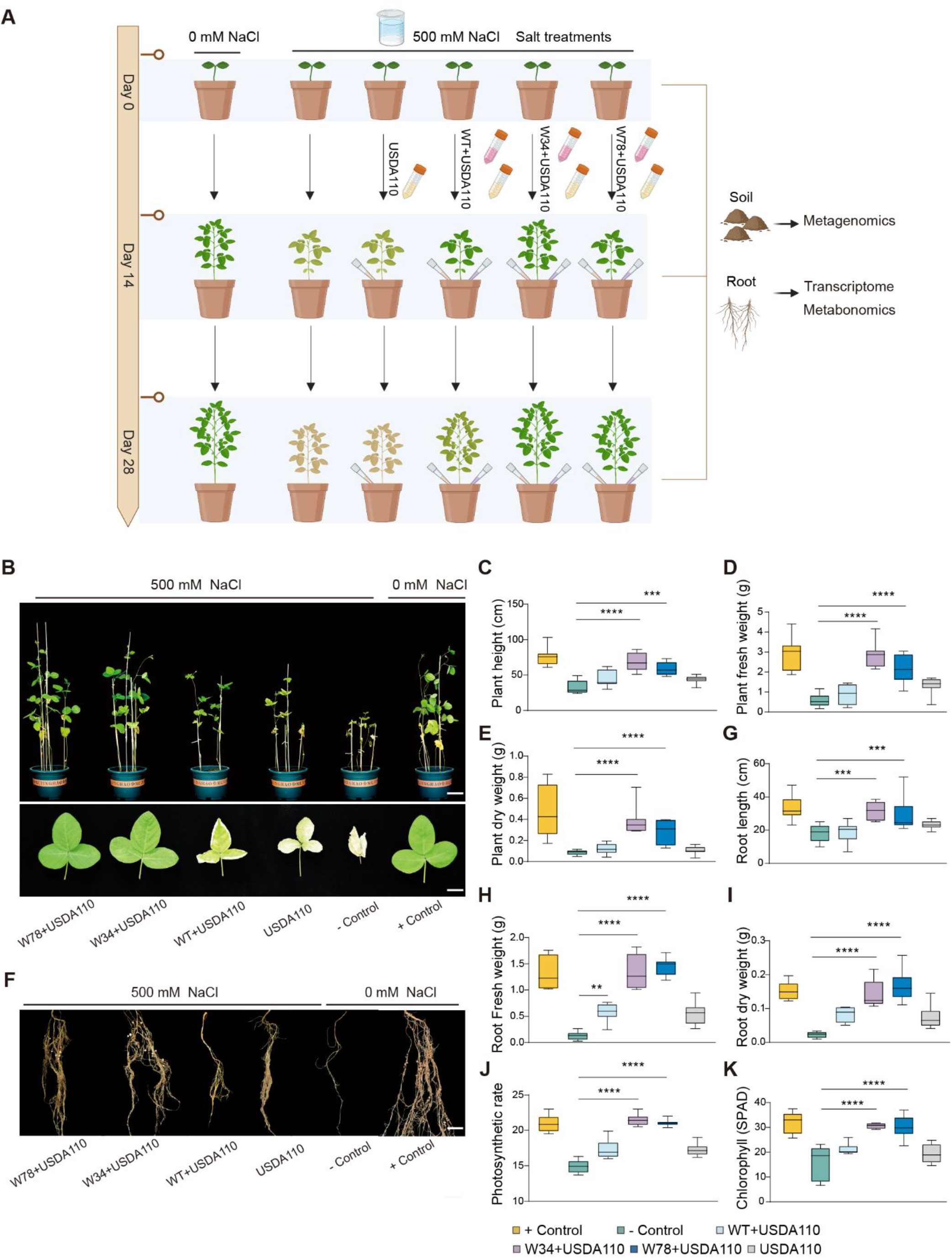
Growth promotion of soybean by ENF *V. natriegens* under saline stress. **(A)** Schematic representation of the experimental design. Soybeans were divided into six treatment groups: (1) positive control group (+Control), normal culture conditions without saline or bacterial inoculation; (2) negative control (−Control), saline-stressed (500 mM NaCl) without bacterial inoculation; (3) *Bradyrhizobium diazoefficiens* USDA110 only; (4) wild type *V. natriegens* WT + USDA110; (5) ENF *V. natriegens* W34 + USDA110; (6) ENF *V. natriegens* W78 + USDA110. Root tissues and rhizosphere soil were collected at 0, 14, and 28 dpi for untargeted metabolomics, transcriptomics, and metagenomic analyses (*n* = 10 biologically independent samples). Created with BioRender.com. **(B)** Soybean shoot phenotype under 500 mM saline stress. Scale bars = 12 cm. **(C, D, E)** Plant height **(C)**, plant fresh weight **(D)**, and dry weight **(E)** under 500 mM saline stress. **(F)** Soybean root phenotype under 500 mM saline stress. Scale bars = 4 cm. **(G, H, I)** Root length **(G)**, root fresh weight **(H)**, and root dry weight **(I)** under 500 mM saline stress. (**J)** Plant photosynthetic rate. (**K)** Plant chlorophyll (SPAD). For boxplot: the center line represents the media, the box bounds represent the lower (Q1) and upper (Q3) quartiles, and the whiskers indicate the minimum and maximum values. Data are presented as mean ± SD (*n* = 10). Error bars represent standard deviations of ten replicates (\**p* < 0.05, \*\**p* < 0.01, \*\*\**p* < 0.001, Student’s *t*-test.

Root system development was also significantly enhanced under saline stress in plants inoculated with ENF *V. natriegens* (Fig. 2F). The W34+USDA110 group exhibited a 90.5-160.9% increase in root fresh weight, a 145.1-229.9 % elevation in dry weight, and a 61.9-126.9% extension in root length compared to WT+USDA110 group (Fig. 2G-I). Additionally, chlorophyll content (SPAD) and photosynthetic rates were significantly enhanced in the ENF *V. natriegens* co-inoculation group (Fig. 2J, K). These results demonstrate that ENF *V. natriegens* mitigates saline stress in soybeans, enhances photosynthetic efficiency and improves soybean productivity.

### ENF V. natriegens enhances soybean nodulation and nitrogenous compound accumulation

Nodule nitrogenase activity, nodule number, and nodule fresh weight are key metrics for assessing symbiotic nitrogen fixation in legumes. Under saline stress, only a very limited number of root nodules were observed in negative controls or soybean inoculated with USDA110 alone. However, soybean inoculated with ENF *V. natriegens* exhibited significantly enhanced nodule formation and nitrogenase activity (Fig. 3A). The W34-inoculated group developed an average of 45 nodules with a total biomass of 1.05 g, representing 10.3-fold increases in nodule number and 8.6-fold enhancements in nodule biomass compared to the WT+USDA110 group. These increases were accompanied by a 5.25-fold elevation in nitrogenase activity (Fig. 3B-D). Paraffin sectioning further confirmed the beneficial effects of ENF *V. natriegens* on nodule growth and biological nitrogen fixation. Nodules from the experimental groups inoculated with ENF *V. natriegens* were deep blue, with round and fully developed infected cells and a uniform distribution of rhizobia, indicative of high metabolic activity. These characteristics were comparable to those observed in the positive control group (+ Control). In contrast, nodules from the WT+USDA110 group exhibited smaller infected cell areas and an uneven distribution of rhizobia (Fig. 3A). These findings suggest that ENF *V. natriegens* enhances soybean growth by promoting nodulation and nitrogenase activity under saline stress.

**Fig. 3.**
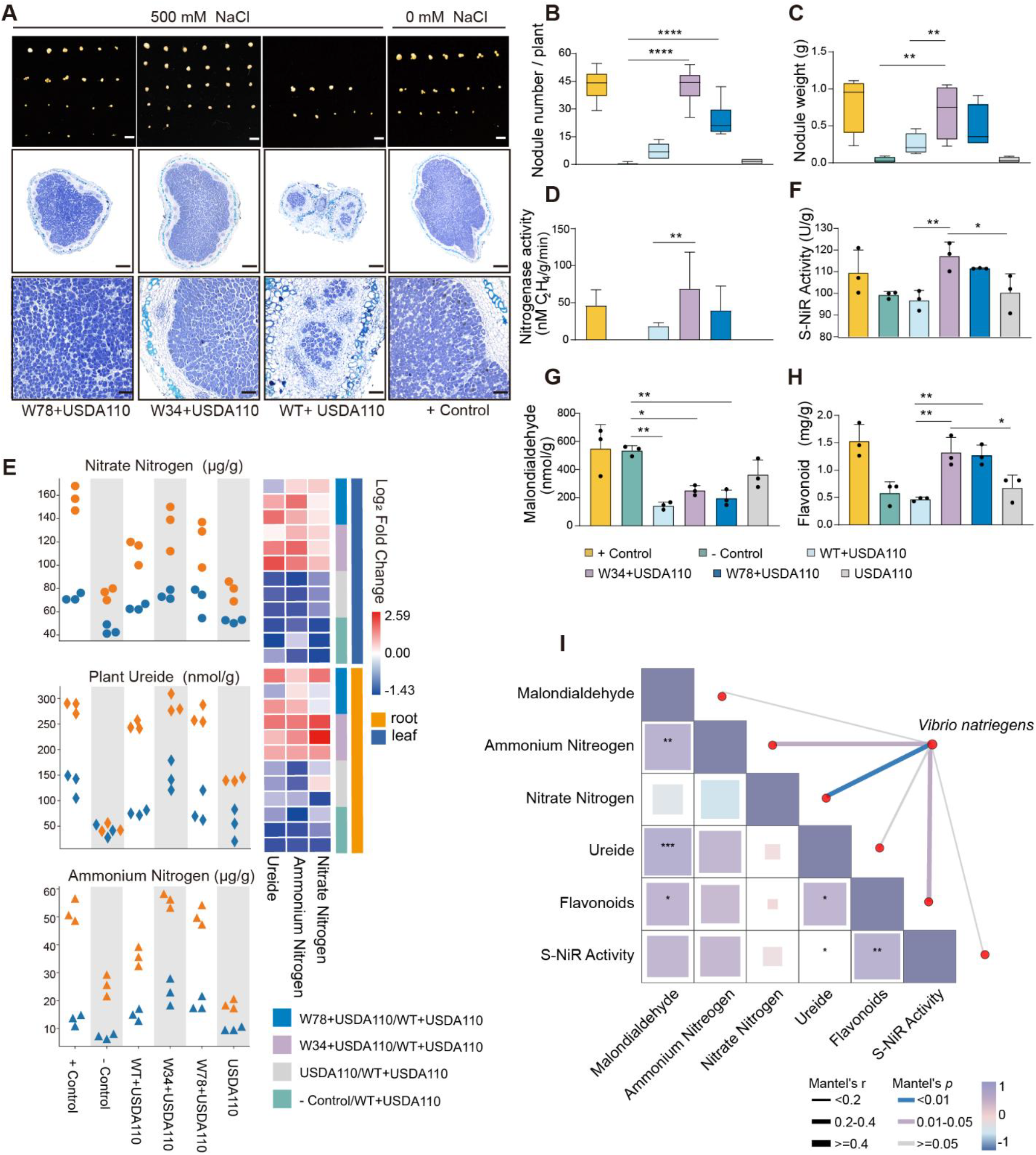
ENF *V. natriegens* enhances soybean nodulation and nitrogenous compound accumulation. (**A)** Effect of different ENF *V. natriegens* inoculations on soybean nodulation phenotypes under 500 mM saline stress. Scale bars = 1 cm. The color of root nodule paraffin sections indicates the presence of living cells within the root nodules. Dark blue staining represents high metabolic activity of rhizobia and well-developed nodules. Scale bars = 500 μm. **(B, C, D)** Nodule number **(B)**, nodule weight **(C)**, and nitrogenase activity **(D)** of soybean under 500 mM saline stress across treatments. Data are presented as mean ± SD (*n* = 10). Statistical significance was determined using Student’s *t*-test (\**p* < 0.05, \*\**p* < 0.01, \*\*\**p* < 0.001). **(E)** Accumulation of nitrogen-containing compounds in the roots and leaves of soybeans under 500 mM saline stress across treatments. **(F)** Nitrate reductase activity in the rhizosphere soil of soybeans under 500 mM saline stress across treatments. Data are presented as mean ± SD (*n* = 3). Statistical significance was determined using Student’s *t-*test (\**p* < 0.05, \*\**p* < 0.01, \*\*\**p* < 0.001). (**G, H)** Malondialdehyde (MDA) concentration (**G**) and flavonoid content (**H**) of soybeans under 500 mM saline stress across treatments. Data are presented as mean ± SD (*n* = 3). Statistical significance was determined using Student’s *t*-test (\**p* < 0.05, \*\**p* < 0.01, \*\*\**p* < 0.001). The boxplot displays the median at the center, with the box bounds representing the lower (Q1) and upper (Q3) quartiles. The whiskers indicate the minimum and maximum values. **(I)** Relationship between ENF *V. natriegens* abundance and nitrogen-containing compounds in soybean, analyzed using *Mantel* tests. Pearson’s correlations were used to assess the relationships between ENF *V. natriegens* abundance and nitrogenous compounds (nitrate nitrogen, ammonium nitrogen, ureide, flavonoids, MDA, and soil nitrate reductase activity). Asterisks indicate the significance levels of correlations. The width of the lines represents the magnitude of the absolute value of *Mantel*’s *r*, while the color of the lines corresponds to the *p*-value significance range (\**p* < 0.05, \*\**p* < 0.01, \*\*\**p* < 0.001).

Nitrogenous compounds, including nitrate nitrogen, ammonium nitrogen, and ureides, play essential roles in soybean nitrogen metabolism by enhancing carbon utilization efficiency and facilitating amino acid assimilation.^22^ As such, their accumulation levels serve as direct indicators of nitrogen fixation capability. Under saline stress, negative control plants exhibited significantly lower nitrogen compound concentrations than those inoculated with ENF *V. natriegens*. The W34+USDA110 group showed a 1.16-to 1.27-fold increase in nitrate nitrogen, a 1.5-to 2.2-fold elevation in ammonium nitrogen, and a 1.96-to 2.38-fold enhancement in ureide content compared to the WT+USDA110 group (Fig. 3E), suggesting that ENF *V. natriegens* contribute to enhanced nitrogen compound accumulation under saline stress.

Soil inorganic nitrogen serves as a key plant-available nutrient, with nitrite reductase activity directly reflecting soil nitrogen transformation efficiency. The highest nitrite reductase activity was observed in the W34+USDA110 group, showing a 1.3-fold increase relative to the USDA110 mono-inoculated group and a 1.08-fold increase compared to the WT+USDA110 group (Fig. 3F), suggesting that W34 exhibits higher nitrogenase activity, which, in turn, enhances soil nitrite reductase activity. Furthermore, soybean plants inoculated with ENF *V. natriegens* exhibited reduced saline-induced cellular damage, as evidenced by increased flavonoid exudation (Fig. 3G) and diminished malondialdehyde (MDA) accumulation (Fig. 3H). *Mantel* test revealed a statistically significant correlation between *V. natriegens* abundance in soil and increased ammonium/nitrate nitrogen levels and flavonoid (*p* < 0.05) (Fig. 3I). These results confirm that ENF *V. natriegens* promotes soybean nitrogen fixation through enhanced nodulation and increased nitrogenous compound accumulation under saline stress.

### ENF V. natriegens activated nitrogen fixation capability of rhizosphere bacteria

To investigate how ENF *V. natriegens* enhances nitrogen fixation in soybeans, we investigated its interaction with soybean roots. GFP-tagged *V. natriegens* maintained a stable rhizosphere population density of 3×10^3^ CFU/g for 10 days post-inoculation (dpi) (Fig. S12). The stereomicroscopic analysis confirmed surface colonization of soybean roots by GFP-tagged *V. natriegens* (Fig. S13). However, confocal laser scanning microscopy detected root autofluorescence in the epidermal layers and vascular bundles (Fig. S14), indicating limited endophytic colonization capacity of *V. natriegens* despite stable rhizosphere persistence under saline stress.

Given its colonization on root surfaces and in saline soils, we hypothesized that *V. natriegens* might influence soybean nitrogen fixation by regulating the rhizosphere bacteria. To test this, we analyzed bacterial community composition at three time points (0 dpi, 14 dpi, and 28 dpi) using Illumina MiSeq sequencing of the V5-V7 region of the 16S rRNA gene. Alpha diversity analysis showed the Chao1 index significantly increased (*p* < 0.01) in the *ENF V. natriegens* groups (Fig. S15A). Principal Coordinate Analysis (PCoA) based on Bray-Curtis dissimilarity matrices further revealed significant shifts in microbial community composition at both 14 dpi (ADONIS: *R*^*2*^ = 0.9795, *p* = 0.001) and 28 dpi (ADONIS: *R*^*2*^ = 0.9727, *p* = 0.001) (Fig. S15B). These findings suggest that ENF *V. natriegens* play a role in shaping the rhizosphere bacteria. To further explore the effect of *V. natriegens* on the rhizosphere bacteria, we analyzed the genus-level compositional profiles. Inoculation with ENF *V. natriegens* led to a significant increase in the relative abundance of nitrogen-fixing bacteria, including *Rhizobium, Bradyrhizobium*, and *Paraburkholderia* (Fig. S15C).

Functional annotation using the *Ncyc* database revealed a progressive increase in nitrogen cycle gene abundance. Rhizosphere soils inoculated with W34 exhibited elevated *nifHDK* gene copy numbers and upregulated expression of genes associated with nitrification, nitrogen fixation, and nitrate reduction pathway (Fig. S15D). Taxonomic annotation of nitrogen-fixation-associated genes in W34-treated rhizosphere soil, based on metagenomic assembled ORFs, showed distinct diversity and distribution patterns. The *nifHDK* genes were predominantly expressed in *Burkholderiales, Hyphomicrobiales, Rhodospirillales* and *Sphingomonadales*, with high annotation proportions at the phylum (96%), class (94%), and order (78%) levels (Fig. S15E). These results demonstrate that ENF *V. natriegens* increases rhizosphere biodiversity and enhances nitrogen fixation efficiency of rhizosphere bacteria.

Next, we hypothesize that ENF *V. natriegens* affects the rhizosphere species composition by stimulating the secretion of metabolites from soybean roots. To identify key metabolite factors in soybeans influenced by ENF *V. natriegens* W34. Weighted Gene Co-expression Network Analysis (WGCNA) was used to integrate differentially accumulated metabolites (DAMs) from soybean roots with rhizosphere bacteria abundance profiles. The relative abundance of *V. natriegens* displayed the highest connectivity with the M-antiquewhite4, which was selected for further analysis (Fig. S16A, B). Using filtering criteria of *R*^*2*^ > 0.6 and *p* < 0.05, key metabolites within this module were identified based on module membership (MM > 0.8) and microbial significance (GS > 0.5) (Fig. S16C). KEGG (Kyoto Encyclopedia of Genes and Genomes) pathway analysis revealed that metabolites associated with *V. natriegens* were enriched in isoflavonoid biosynthesis, phenylpropanoid biosynthesis, flavone and flavonol biosynthesis, and flavonoid biosynthesis pathways (Fig. S16D). *Paraburholderia*, a nitrogen-fixing bacteria with increased relative abundance in ENF *V. natriegens*-treated rhizosphere soil, exhibited significant positive correlations (*p* < 0.05) with coniferyl aldehyde (MN9636), trans-cinnamate (MP2946), and apigenin (MN882) (Fig. S17). These finding suggest that ENF *V. natriegens* enhances the abundance of rhizosphere nitrogen-fixing bacteria by stimulating flavonoid metabolites secretion from soybean roots.

## DISCUSSION

Nitrogen is a key element for microbial and plant growth, thereby mediating plant resistance to abiotic stress.^28^ In this study, we demonstrated that ENF *V. natriegens* carrying the *nif* gene cluster from *A. vinelandii* DJ enhanced rhizosphere nitrogen fixation by stimulating flavonoid secretion. Concomitantly, ENF *V. natriegens* upregulated neohesperidin biosynthesis in soybean roots, which modulated the expression of nitrogen metabolism genes and symbiotic nitrogen fixation genes, ultimately boosting soybean nitrogen-fixing capacity under saline stress.

Naturally occurring nitrogen-fixing genes are typically clustered, with sizes ranging from 13 kb in *P. polymyxa* WLY78 to 65 kb in *P. stutzeri* A1501. Larger gene clusters tend to encode bioactive nitrogenase, electron transport chains, and oxygen protection mechanisms,^29^ allowing for more effective nitrogen fixation under diverse conditions. However, transferring *nif* gene clusters across species using inducible regulatory systems presents challenges in real-world applications due to safety concerns, unpredictable induction timing, and unstable nitrogenase expression.^30,31^ To overcome these limitations, four *nif* gene clusters with high heterologous nitrogenase activity in *E. coli* were selected: those from *P. polymyxa* WLY78, *K. oxytoca* M5al, *A. vinelandii* DJ, and *P. stutzeri* A1501.^32^ Compared to *P. polymyxa* WLY78 and *K. oxytoca* M5al, *A. vinelandii* DJ contain longer *nif* clusters (65 kb), which may contribute to their enhanced functionality (Fig. 1A). Although all four clusters conferred nitrogen fixation capability, the W34 carrying *A. vinelandii* DJ gene clusters, demonstrated superior growth rates and nitrogenase activity (Fig. 1B, C), which was consistent with previous research.^32^ As obligate aerobes that fix nitrogen under ambient conditions, *A. vinelandii* DJ demonstrate improved oxygen resilience, facilitating nitrogen fixation in soil environments.^33,34^

Notably, the *A. vinelandii* DJ gene cluster enhances the expression of upstream glutamate metabolism genes (*gltBD, gdh, glnA*) in ENF *V. natriegens* W34(Fig. 1E). Glutamine synthetase (GS) and glutamate synthase (GOGAT) constitute the core pathway of microbial nitrogen assimilation. Improving the efficiency of the GS/GOGAT cycle is considered an effective approach to enhance nitrogen utilization efficiency,^35,36^ thereby supplying carbon skeletons for the TCA cycle and maintaining the carbon-nitrogen balance in the bacteria.^37^ Since GS is a key enzyme for ammonium assimilation in soybean, its overexpression is known to enhance plant nitrogen use efficiency.^38,39^ Inoculation with ENF *V. natriegens* W34 significantly upregulated glutamine synthetase (GS) genes (*SoyZH13_14G150200, SoyZH13_19G060300*) in soybean, with expression levels exceeding those in other treatment groups at both 14 and 28 dpi (Fig. S18). Consistent with these findings, the relative abundance of ENF *V. natriegens* showed a positive correlation with *SoyZH13_06G10900* (K10534: nitrate reductase) and *SoyZH13_08G267800* (K02575: nitrate transporter) (Fig. S19), suggesting that ENF *V. natriegens* plays a critical role in enhancing soybean nitrogen metabolism efficiency.

Synthetic biology presents novel solutions for sustainable agriculture. In this study, ENF *V. natriegens* maintained its growth-promoting effects under 500 mM NaCl stress (Fig. S20), whereas naturally saline-tolerant PGPBs and synthetic microbial communities (SynComs) exhibited significant growth-promoting effects only at NaCl concentrations ≤ 200 mM (table. S4). Consistent results were obtained from peanut pot trials under 500 mM NaCl stress (Fig. S21), demonstrating the broad applicability of this approach in sustainable agriculture. *LefSe* analysis identified microbial species differences across treatments.^40^ At 14 dpi, major species shifts were observed between the positive and negative control groups, likely due to saline-induced changes in the rhizosphere community. However, species differences expanded across all treatment groups, with *Vibrio, Soliminas*, and *Verrucomicrobiaceae* dominating in the W34+USDA110 group at 28 dpi (Fig. S22). This suggests that W34 is a broad-host-range nitrogen-fixing bacteria and establishes a stable presence in the rhizosphere.

This study establishes a synthetic biology framework for engineering next-generation multifunctional plant growth-promoting bacteria (PGPB). It also provides insights into how ENF *V. natriegens* enhances soybean growth and nitrogen fixation under saline stress. These discoveries serve as a foundation for advancing precision agriculture through synthetic biology-driven microbial engineering, offering a sustainable and transformative approach to improving crop stress resilience in saline farmland ecosystems.

## Funding

National Key Research and Development Program (China) 2021YFA0909500

National Natural Science Foundation (China) 32030004

National Natural Science Foundation (China) 32370106

National Natural Science Foundation (China) U22A20444

Shanghai Municipal Science and Technology Major Project

## Author contributions

Conceptualization: Hongzhi Tang, Wanjing Wu Methodology: Wanjing Wu

Investigation: Wanjing Wu

Visualization: Wanjing Wu Funding acquisition: Hongzhi Tang,

Project administration: Hongzhi Tang Supervision: Hongzhi Tang

Writing – original draft: Wanjing Wu Writing – review & editing: Wanjing Wu

## Data and materials availability

The clean reads of RNA-seq in this paper have been deposited in the SRA database (SRA Accession No.: PRJNA1255886, PRJNA1256108, PRJNA1256128). The raw metabolomics data reported in this paper have been deposited in the MTBLS, MetaboLights (https://www.ebi.ac.uk/metabolights/; Accession No.: REQ20250427210173). All other data are included in the manuscript and/or supporting information.

## Supplementary Materials

Materials and Methods Figures. S1 to S28

Tables. S1 to S7

Table S2. Excel file containing additional data too large to fit in a PDF

Table S3. Excel file containing additional data too large to fit in a PDF

References (*52-80*)

