## Supplemental figure for "Breaking the Salinity-Nitrogen Fixation Trade-Off: Engineering A Synthetic Nitrogen-Fixing *Vibrio natriegens* Strain"

Wanjing Wu[1]†, Haiyang Hu[1]†, Jinzhong Tian[2]†, Weiwei Wang[1]*, Ling Li[3], Ping Xu[1]*, Junbiao Dai[3], Hongzhi Tang[1]*

*Corresponding author: H. Z. Tang or W. W. Wang or P. Xu

**The PDF file includes:**

Materials and Methods

Figures. S1 to S28

Tables. S1 to S7

Table S2. Excel file containing additional data too large to fit in a PDF

Table S3. Excel file containing additional data too large to fit in a PDF

References (1-29)

### Materials and Methods

#### *Bacterial strains and growth media*

All bacterial strains and their derivatives used in this study are listed in table. S5. *E. coli* DH10α was used for cloning. Bacteria were grown in rich media, including LB medium, which contained 10 g $L^{-1}$ tryptone, 5 g $L^{-1}$ yeast extract, and 10 g $L^{-1}$ NaCl. LBV2 medium was composed of 10 g $L^{-1}$ tryptone, 5 g $L^{-1}$ yeast extract, 0.3 g $L^{-1}$ KCl, 5 g $L^{-1}$ $MgCl\cdot 6H_2O$, and 22 g $L^{-1}$ NaCl. TY medium contained 5 g $L^{-1}$ tryptone, 3 g $L^{-1}$ yeast extract, and 0.7 g $L^{-1}$ $CaCl_2\cdot 2H_2O$. Minimal media included M9 medium, which consisted of 0.25 g $L^{-1}$ $MgSO_4\cdot 7H_2O$, 25 g $L^{-1}$ NaCl, 0.1 g $L^{-1}$ $CaCl_2\cdot 2H_2O$, 64 g $L^{-1}$ $Na_2HPO_4\cdot 7H_2O$, 15 g $L^{-1}$ $KH_2PO_4$, 5 g $L^{-1}$ $NH_4Cl$, and 20 g $L^{-1}$ glucose, adjusted to pH 7.4. A trace element solution and a vitamin solution were also added to the medium. The trace element solution contained $Na_2$-EDTA (0.5 g $L^{-1}$), $H_3BO_3$ (2.86 g $L^{-1}$), $MnCl\cdot 4H_2O$ (1.81 g $L^{-1}$), $ZnSO_4\cdot 7H_2O$ (0.222 g $L^{-1}$), $Na_2MoO_4\cdot 2H_2O$ (0.39 g $L^{-1}$), $CuSO_4\cdot 5H_2O$ (0.079 g $L^{-1}$), and Co $(NO_3)_2\cdot 6H_2O$ (0.0494 g $L^{-1}$). The vitamin solution contained riboflavin (0.5 g $L^{-1}$), p-aminobenzoic acid (0.08 g $L^{-1}$), nicotinic acid (0.5 g $L^{-1}$), biotin (0.12 g $L^{-1}$), thiamine HCl (0.8 g $L^{-1}$), calcium pantothenate (0.8 g $L^{-1}$), inositol (0.48 g $L^{-1}$), and cobalamine (0.001 g $L^{-1}$). The glucose, phosphate, and trace element solutions were autoclaved separately, while the vitamin solution was filter-sterilized. Nitrogen-free medium was prepared by removing the nitrogen source from minimal media.[1] Cells were grown in N-depression media for nitrogenase activity assays. While not strictly nitrogen-free, N-depression media contained 1.43 mM serine to promote ribosomal RNA production and accelerate nitrogenase biosynthesis. Antibiotics were used at the following concentrations: kanamycin (50 μg $mL^{-1}$), spectinomycin (100 μg $mL^{-1}$), tetracycline (15 μg $mL^{-1}$), gentamicin (15 μg $mL^{-1}$) for *E. coli*, while *c*hloramphenicol (11 μg $mL^{-1}$) was used for *V. natriegens*.

#### *Construction of nif clusters*

To construct large *nif* clusters on mobilizable plasmids, DNA sequence information from *P. polymyxa* WLY78, *K. oxytoca* M5al, and *A. vinelandii* DJ was obtained from NCBI. The *nif* cluster DNA was synthesized *de novo* in separate fragments using gene synthesis (Genomics Institution) and used as templates for PCR amplification and assembly. Genomic DNA from *P. stutzeri* A1501 was purified using a *Bocai* DNA purification kit (K132) following the isolation protocol for Gram-negative bacteria. This DNA was also used as a template for PCR amplification and assembly. Each *nif* cluster was amplified into several fragments ranging from 4–5 kb, with 150 bp upstream and downstream linkers flanking the 5′ and 3′ ends. The fragments were assembled into the linearized *E. coli*–yeast shuttle vectors pRS415 for *E. coli* and *V. natriegen*s using yeast recombineering.[2] The amplified DNA fragments were assembled into a single large plasmid via a one-pot yeast assembly procedure. Once assembled, the *nif* cluster plasmids were isolated from yeast using a Zymoprep yeast miniprep kit (Zymo Research, cat# D2004) and subsequently transformed into *E. coli.* The purified plasmid was extracted from *E. coli* and sequenced to verify correct assembly and sequence integrity. *E. coli* strains containing mutation-free plasmid were stored for further experiments.

#### *Construction of nitrogen fixation in V. natriegens*

To obtain large *nif* clusters from a mutation-free plasmid in *E. coli,* nif cluster plasmids were extracted using the Macherey-Nagel NucleoBond BAC 100 kit (740579). The nitrogen fixation gene fragments, flanked by homologous arms of *V. natriegens, were isolated* through enzymatic digestion and subsequently integrated into chromosome 2 of *V. natriegens* via natural transformation (*15*). For natural transformation, *V. natriegens* were induced by overnight growth (15 hours) in LBV2 supplemented with 0.1 mM IPTG at 30 °C in an orbital shaker incubator. A 3.5 μL aliquot of the overnight culture was directly diluted into 350 μL of Instant Ocean Medium (IOM, 28 g $L^{-1}$) supplemented with 0.1 mM IPTG. Approximately 3 μg of the transforming DNA fragment was then added, and the reactions were incubated statically at 30 °C for six hours. Following this incubation, 1 mL of LBV2 was added, and the cultures underwent outgrowth at 30 °C with shaking (220 rpm) for two hours. The resulting cultures were then spread onto antibiotic LBV2 agar plates to select for successful integration of the desired gene fragments.

***Growth curves measurement***

Engineered and wild type *V. natriegens* were incubated overnight in LBV2 medium, washed three times with nitrogen-free medium, and subsequently cultured in nitrogen-free medium with an initial $OD_{600}$ of 0.01. The strains were maintained under nitrogen-free conditions in an anaerobic incubator (90% $N_2$, 5% $CO_2$ and 5% $H_2$), $OD_{600}$ measurements were taken at regular intervals following removal from the incubator.

***Nitrogenase assays***

Cultures were initiated by inoculating a single colony into 5 mL of LBV2 supplemented with the appropriate antibiotics in sterile tubes and incubated overnight at 30 °C and 220 rpm in a Multitron incubator. The bacterial solution was then washed twice with M9 medium, and 50 μL aliquots of the overnight cultures were diluted in 5 mL of M9 medium containing 5 g $L^{-1}$ $NH_4Cl$. The cultures were incubated overnight at 30 °C and 220 rpm. After washing twice with nitrogen-free medium, the cultures were diluted to an $OD_{600}$ of 0.4 in 14 mL of N-depression medium and supplemented with the appropriate antibiotics and 1.43 mM serine to facilitate nitrogenase depression. These cultures were transferred into 20-mL headspace glass vials with PTFE-silicone septa screw caps. The headspace of the vials was replaced with 100% argon gas using a vacuum manifold. Acetylene, freshly generated from $CaC_2$ in a Burris bottle, was injected into each vial to a final concentration of 10% (v/v) to initiate the reaction. Acetylene reduction was carried out for 20 hours at 30 °C with shaking at 220 rpm in an Innova shaking incubator (New Brunswick) to prevent cell aggregation. The reaction was quenched by adding 0.5 mL of 4 M NaOH to each vial. To quantify ethylene production, culture headspace was withdrawn using a gas-tight syringe and manually injected into a SHIMADZU GC2030 gas chromatograph. Ethylene production was determined by integrating the area under the peak using ChemStation software, and the corresponding formula was applied for calculation.

***Assessment of the growth-promoting ability of V. natriegens***

*Determination of indole-3-acetic acid (IAA) production by V. natriegens*

Following activation, the test strain was inoculated into Landy medium supplemented with 3 mM L-tryptophan at an inoculation volume of 0.1%. The culture was incubated at 25 °C and 140 rpm in the dark for 72 hours. A 1 mL aliquot of the culture was then centrifuged at 12,000×*g* for 5 minutes. The resulting supernatant (1 mL) was mixed with an equal volume of Salkowski reagent and incubated at room temperature in the dark for 30 minutes. Optical density measurements were recorded at 530 nm, using blank medium as the control. IAA production (mg $L^{-1}$) was determined based on a standard curve prepared with pure IAA. For the IAA standard curve, solutions of IAA at concentrations of 0, 10, 20, 25, 40, and 50 mg $L^{-1}$ were used as reference standards. Landy Medium Composition (per liter): 20 g glucose, 1 g yeast extract, 5 g

$L^{-1}$ glutamic acid, 2 mg L-phenylalanine, 1 g L-tryptophan, 0.5 g KCl, 1 g $KH_2PO_4$, 0.5 g $MgSO_4 \cdot 7H_2O$, 5 mg $MnSO_4 \cdot 4H_2O$, 0.16 mg $CuSO_4 \cdot 7H_2O$, 0.15 mg $FeSO_4 \cdot 7H_2O$, with distilled water adjusted to 1000 mL, pH 7.0. Salkowski Reagent: 10.8 M $H_2SO_4$ containing 4.5 g $FeCl_3$.

***Determination method of siderophore production by V. natriegens.***

The test strain was first inoculated into LBV2 mediums and cultured overnight. A 1% inoculum was then transferred into 5 mL of LBV2 liquid medium and incubated at 30 °C with shaking at 220 rpm overnight. Optical density (OD) measurements were taken to ensure uniform OD values across all groups. The culture was then centrifuged at 5,000 rpm for 10 minutes, and 5 mL of the resulting supernatant was filtered to remove bacterial cells. The supernatant was aliquoted into three sterile 15 mL centrifuge tubes, with 1 mL in each tube. One milliliter of CAS assay solution was added to each tube, and the reaction was allowed to proceed. The OD at 630 nm ($OD_{630}$) was measured to determine the absorbance value ($A_s$). Double-distilled water served as the blank control. A mixture of uninoculated medium and CAS assay solution in equal volumes was used to measure the $OD_{630}$, yielding the absorbance value ($A_r$). The $A_s/A_r$ ratio represented the relative siderophore content in the samples, with each measurement repeated three times. The $A_s/A_r$ ratio ranges from 1.0 to 0, with intervals of 0.2. Each decrease of 0.2 was assigned one "+" to indicate siderophore production. Strains with high siderophore production typically exhibited an $A_s/A_r$ ratio below 0.5.

- CAS Blue Assay Solution:
  - Solution A: 60.5 mg CAS dissolved in 50 mL deionized water, with 10 mL $FeCl_3$ solution (containing 1 mM $FeCl_3 \cdot 6H_2O$, 10 mM HCl) added.
  - Solution B: 72.9 mg HDTMA (hexadecyltrimethylammonium bromide) dissolved in 40 mL deionized water.

Solution A was slowly poured into Solution B along the inner wall of a beaker to obtain the CAS blue assay solution.

***RNA-seq analysis***

Different engineered strains of *V. natriegens* were cultured overnight in LBV2 medium. Following centrifugation at 6000 × *g* for 20 minutes at 4 °C, the cells were washed twice with nitrogen-free medium to completely remove residual carbon and nitrogen sources, as well as secondary metabolites. Each engineered strain was then inoculated into one nitrogen-free medium and two M9 media, with an initial OD of 0.01. The cultures were incubated under different growth conditions: aerobic with a nitrogen source, anaerobic with a nitrogen source, and anaerobic without a nitrogen source. Each engineered strain was tested in triplicate for each growth condition. After a defined growth period, the cells were harvested by centrifugation at 6000 × *g* for 20 minutes at 4 °C and rapidly frozen in liquid nitrogen. RNA-seq was conducted by Shanghai Personalbio Technology Co. Ltd., China. Total RNA was extracted using TRIzol® reagent (R0016, Beyotime, China), and genomic DNA was removed with DNase I (TaKara, Japan). Ribosomal RNA (16S and 23S rRNA) was depleted from the total RNA using a Ribo-Zero Magnetic kit (Epicenter Biotechnologies, WI, USA). mRNA was subsequently fragmented (~200 bp) using a fragmentation buffer. Complementary DNA (cDNA) synthesis was performed through reverse transcription using the SuperScript double-stranded cDNA synthesis kit (Invitrogen, CA) with random hexamer primers (Illumina, USA). During second-strand cDNA synthesis, deoxyuridine triphosphate (dUTP) was incorporated in place of deoxythymidine triphosphate (dTTP) to generate blunt-ended cDNA fragments. These double-stranded cDNA fragments then underwent end repair, phosphorylation, 3′ adenylation, and adapter ligation. The second-strand cDNA containing dUTP was selectively degraded using uracil-N-glycosylase (UNG) enzyme. Following degradation, cDNA fragments were separated on a 2% agarose gel, and DNA fragments of approximately 200 bp were extracted for cDNA library construction. PCR amplification of the cDNA libraries was performed using Phusion DNA polymerase (NEB, USA) for 15 cycles. The libraries were quantified using a microfluorometer (TBS-380, TurnerBioSystems, USA) and sequenced on an Illumina HiSeq× Ten platform using paired-end sequencing. Bioinformatics analyses were conducted using the cloud-based platform of Shanghai Personalbio Technology Co. Ltd., based on sequencing data generated by the Illumina platform.

***RNA extraction and quantitative real-time PCR analysis***

The plants were treated or not treated with different neohesperidin for 14 days. Uninoculated and inoculated roots were collected and total RNA was extracted using the TRIzol reagent according to the manufacturer's protocol (Transgen, Beijing, China). For mRNA quantification, the first-strand complementary DNA (cDNA) synthesis was performed with the Transcriptor First Strand cDNA Synthesis Kit instructions (Transgen, Beijing, China). qRT–PCR was performed on 3 biological replicates using the Bio-Rad CFX96 real-time PCR detection system with SYBR (Transgen, Beijing, China). *GmActin11* expression was used as an internal control. Data were collected using the corresponding real-time PCR detection systems.The relative transcript abundance was determined by calculating the ratio of the target gene expression value of the target gene to that of *GmActin11* using the $2^{-\Delta\Delta CT}$ (threshold cycle) method. Primers are listed in table. S6

***Plant material and pot experiments***

Soybean (*Glycine* max cv. Zhonghuang 13) and the rhizobial species *Bradyrhizobium diazoefficiens* USDA110 were used in this study. Nodule bacteria medium (TY) and LBV2 were used to enrich the rhizobia and *V. natriegens*, and detailed medium formulations were provided in the "Bacterial strains and growth media" section. To prepare bacterial inoculants, B. diazoefficiens USDA110 was inoculated into TY liquid medium at a 1% volume and incubated at 28 °C for 48 hours. The culture was then centrifuged at 5,000 rpm for 20 minutes to collect viable bacteria, which were resuspended in sterile water to an $OD_{600}$ = 0.08. Similarly, *V. natriegens* was inoculated into an LBV2 liquid medium at a 0.1% volume and incubated at 30 °C for 10 hours. The culture was centrifuged at 5,000 rpm for 20 minutes, and the bacteria were resuspended in sterile water containing 500 mM NaCl to an $OD_{600}$ of 3. Additionally, a low-nitrogen nutrient solution containing 0.25 mM $NO_3^-$ was prepared. The specific composition of the nutrient solution is shown in table S7.

To characterize the effects of *V. natriegens* and USDA110 on soybean-associated communities and functions in saline soil environments, wild soybean plants were grown in agricultural soil under controlled conditions. Seeds were surface-sterilized with 0.15% mercuric chloride for 20 minutes and washed six times with sterile water. They were then germinated in Petri dishes with sterile water inside a growth chamber. After germination, seedlings were transferred to plastic pots containing 200 g of field soil and incubated at 26 °C on a 16-hour/8-hour light/dark cycle. After 10 days of growth, uniformly developed seedlings were transplanted into plastic pots containing 2 kg of nutrient-rich soil. Based on preliminary experiments, different NaCl concentrations were tested, and 500 mM NaCl was determined to be optimal. Once the transplanted soybean seedlings stabilized in the soil, they were treated with a low-nitrogen nutrient solution containing 500 mM NaCl, while a low-nitrogen nutrient solution without NaCl served as the control. The experiment was conducted with four biological replicates, where ten wild soybean plants per pot constituted one replicate.

To assess dynamic changes in the root-associated microbiome and rhizosphere metabolome, rhizosphere soil samples and soybean root tissue samples were collected at three time points: day 0, day 14, and day 28 after saline stress. The soil samples were stored at -80 °C, and root tissue samples were cleaned before storage at -80 °C.

***Physiological indicators and biological nitrogen fixation***

Plant samples were collected at different growth stages. Harvested soybeans were cut at the root-stem junction, and the aboveground and belowground parts were separated to measure fresh weight, plant height, and root length. The samples were then dried in an oven at 65 °C for 48 hours to determine their dry weight and water content.

Following the method described by root samples from eight plants were collected, and excess soil was gently shaken off.[3] Nodules were weighed and recorded. The samples were then placed in 60 mL serum bottles to create a closed system. Using a syringe, 3.0 mL of air was extracted from each bottle and immediately replaced with 3.0 mL of acetylene gas. The bottles were incubated at 28 °C for 2 hours. Following incubation, 5.0 mL of gas was extracted and injected into a 20 mL headspace vial. Ethylene peak area measurements were obtained via gas chromatography.

*Preparation of toluidine blue-stained paraffin sections*

To observe nodule morphology, nodules from different treatment groups were collected and fixed in formalin-acetone alcohol (FAA) fixative solution for the preparation of toluidine blue-stained paraffin sections. The stained sections were then examined under an optical microscope. Toluidine blue, a synthetic dye, selectively stains living cells. In paraffin sections of root nodules, darker staining indicates a higher abundance of viable cells, primarily rhizobia.

***Measurement of physiological and biochemical indicators in soybean and soil***

To assess the effects of *V. natriegens* on plant nitrogen-containing compounds (nitrate nitrogen, ammonium nitrogen, and urea), malondialdehyde, flavonoids, and soil nitrite reductase activity, plant tissue samples were rapidly frozen in liquid nitrogen and ground into a fine powder. Equal amounts of tissue were weighed and analyzed using corresponding assay kits (Solarbio).

***Measurement of chlorophyll concentration***

Chlorophyll concentration was measured as described previously by *Zhang* et al.[4] Briefly, approximately 0.2 g of leaves were cut into 2 mm pieces and placed in a tube, followed by the addition of 4 mL of acetone–ethanol (1:1) extraction solution. The tubes were incubated in the dark for 14 hours with shaking to ensure thorough extraction. Once the leaves turned white, the solution was diluted to 10 mL using the extraction solvent. The absorbance of the extract was measured at 645 nm and 663 nm using a microplate reader (Thermo Fisher Scientific). The chlorophyll concentration was calculated using the following formula: Chlorophyll concentration $(\mathrm{mg\ g^{-1}}) = (8.02\times A_{663}+20.21\times A_{645})\times V/(1000\times W)$.

where: A663 and A645 are the absorbance values at 663 nm and 645 nm, respectively; V is the total volume of the extract (mL), and W is the fresh weight of the leaf sample (g).

***Determination of bacterial colonization***

To evaluate the colonization ability of *V. natriegens* in soil, a green fluorescent protein (GFP)-labeled strain was constructed. In a pot experiment, soil samples were collected every 7 days and incubated in M9 liquid medium on a shaker at 30 °C for 2 hours. The supernatant was then diluted to $10^6$ and plated on LBV2 agar containing appropriate resistance markers. After overnight incubation at 30 °C, colonies were verified using PCR, and fluorescence microscopy was used to confirm the presence of the GFP-labeled strain.

For microscopic imaging, a confocal laser scanning microscope was used to visualize bacterial colonization. Roots inoculated with the fluorescent *V. natriegens* strain were manually sectioned and stained with 20 μg/mL propidium iodide (PI) before imaging. A spinning disk confocal microscope was used to capture images, and bacterial colonization was assessed in twenty soybean plants.

***Transcriptome analysis***

To investigate the effect of sodium-demanding *V. natriegens* on the transcriptome of soybean roots, plant root materials were collected at different growth periods. The roots were washed thoroughly, sonicated, and frozen after the washing buffer was removed. Six biological replicates were prepared for each condition Transcriptome assays were conducted by Shanghai Personalbio Technology Co. Ltd., China. Total RNA was extracted from root tissue using TRIzol® Reagent following the manufacturer's instructions, and genomic DNA was removed using DNase I. RNA quality was assessed with a 2100 Bioanalyzer (Agilent) and quantified using an ND-2000 spectrophotometer (NanoDrop Technologies). RNA-seq transcriptome libraries were prepared using the TruSeq™ RNA Sample Preparation Kit (Illumina, San Diego, CA) with 1 μg of total RNA. Briefly, poly(A)+ messenger RNA was isolated using oligo (dT) beads and fragmented using a fragmentation buffer. Double-stranded cDNA was synthesized using a SuperScript double-stranded cDNA synthesis kit (Invitrogen, CA) with random hexamer primers (Illumina). The synthesized cDNA underwent end repair, phosphorylation, and A-tailing following Illumina's library construction protocol. Libraries were size-selected for 300 bp cDNA target fragments using 2% Low Range Ultra Agarose, followed by PCR amplification using Phusion DNA polymerase (NEB) for 15 cycles. After quantification using a TBS380 fluorometer, the paired-end RNAseq library was sequenced on an Illumina NovaSeq 6000 platform (2 × 150 bp read length). Raw paired-end reads were trimmed and quality-controlled using fastp (version0.19.5, https://github.com/OpenGene/fastp) with default parameters. Clean reads were aligned to the reference genome in orientation mode using HISAT2 (version2.1.0, http://ccb.jhu.edu/software/hisat2/index. shtml). Gene expression levels were quantified using the transcripts per million reads (TPM) method using RSEM. Differentially expressed genes were identified based on a false discovery rate (FDR) $\leq 0.001$ together and an absolute value of $\log_2$(relative expression level) $\geq 1$.

***Metabolomic analyses***

To identify key rhizosphere metabolites in responses to stress and *V. natriegens*, an untargeted metabolomics approach was employed to analyze root metabolites. The metabolomic assays were conducted by Shanghai Personalbio Technology Co. Ltd., China. For untargeted metabolomics analysis, root material was collected at different growth stages, with six biological replicates per condition. Metabolites were extracted using a pre-cooled acetonitrile: methanol:

$H_2O$ (2:2:1, v/v/v) solvent system. This solvent combination was selected to enhance the extraction efficiency of metabolites with varying polarities, facilitate cell membrane penetration, preserve metabolite stability, and ensure compatibility with mass spectrometry for high-quality spectral data acquisition. Following centrifugation at 10,000 × *g* for 10 minutes, the supernatant was collected for LC-MS/MS analysis. Quality control (QC) samples were prepared by pooling equal aliquots from all collected samples. One QC sample was injected before analysis and after every ten runs to monitor analytical variance.

Untargeted metabolomics was performed using a Vanquish UHPLC system coupled with a Q Exactive HF-X mass spectrometer. Bioinformatics analysis was conducted on a cloud platform provided by Shanghai Personalbio Technology Co. Ltd., based on the data generated by the Illumina platform.

***Metagenomic sequencing***

To investigate the effects of *V. natriegens* addition on rhizosphere soils, metagenomic assays were conducted by Shanghai Personalbio Technology Co. Ltd., China. Rhizosphere soil samples were collected at different growth stages and subjected to metagenomic shotgun sequencing. Libraries were prepared without an amplification step, and sequencing was performed on the Illumina HiSeq X-Ten platform using 2 × 150 bp paired-end reads at Majorbio Bio-Pharm Technology. Reads containing adapters, low-quality bases, or more than 10% undefined bases were removed. A total of ~264 GB of clean data was obtained from 16 samples. High-quality reads were assembled using Megahit software, and contigs shorter than 300 bp were discarded. Gene prediction was performed using MetaGene, followed by clustering with CD-Hit at 90% identity and 90% coverage to remove redundant sequences. For taxonomic and functional analyses, genes were aligned (BLASTp) against the NCBI-nr, KEGG, and COG databases using DIAMOND with an E-value cutoff of $10^{-5}$. Bioinformatics analysis was conducted using a cloud platform (Shanghai Personalbio Technology Co. Ltd.) based on the data generated by the Illumina platform.

***WGCNA analysis***

Weighted gene co-expression network analysis (WGCNA) is an unsupervised clustering tool used to identify highly interconnected gene clusters in different modules across samples. In this study, plant root metabolomes and rhizosphere soil macro genomes were jointly analyzed using WGCNA to identify key metabolites and genes. This approach was used to correlate gene expression patterns with plant root metabolites and determine the effects of *V. natriegens* addition on soil macrogenomes and root metabolites. The following parameters were applied: TOMType = unsigned, minModuleSize = 30, mergeCutHeight = 0.35, and default values for all other parameters.

***Biological function and gene set enrichment analyses***

To explore the biological functions of differentiation-related modules, enrichment analyses were performed, including GO term and KEGG pathway enrichment analyses. GO enrichment analysis was conducted using GOATOOLS (https://github.com/tanghaibao/goatools), whereas KEGG pathway enrichment analysis was performed using KOBAS.

***Statistical analysis***

Primary data processing and organization were performed in Microsoft Excel (2021). Statistical analyses were conducted in GraphPad Prism 8.

For comparisons between two groups, an unpaired two-tailed *t*-test was used. For comparisons among more than two groups, one-way ANOVA was applied. Depending on the research question, one-way ANOVA was followed either by no post hoc test or by Tukey's post hoc test (for pairwise comparisons among all groups), as indicated in the figure legends. Data are expressed as means, with error bars representing standard deviation (SD) unless specified otherwise. A detailed description of the statistical analyses is provided in the figure legends. Statistical significance was defined as follows: *$p < 0.05$, **$p < 0.01$, ***$p < 0.001$, ****$p < 0.0001$. ns, not significant

**Fig. S1.**

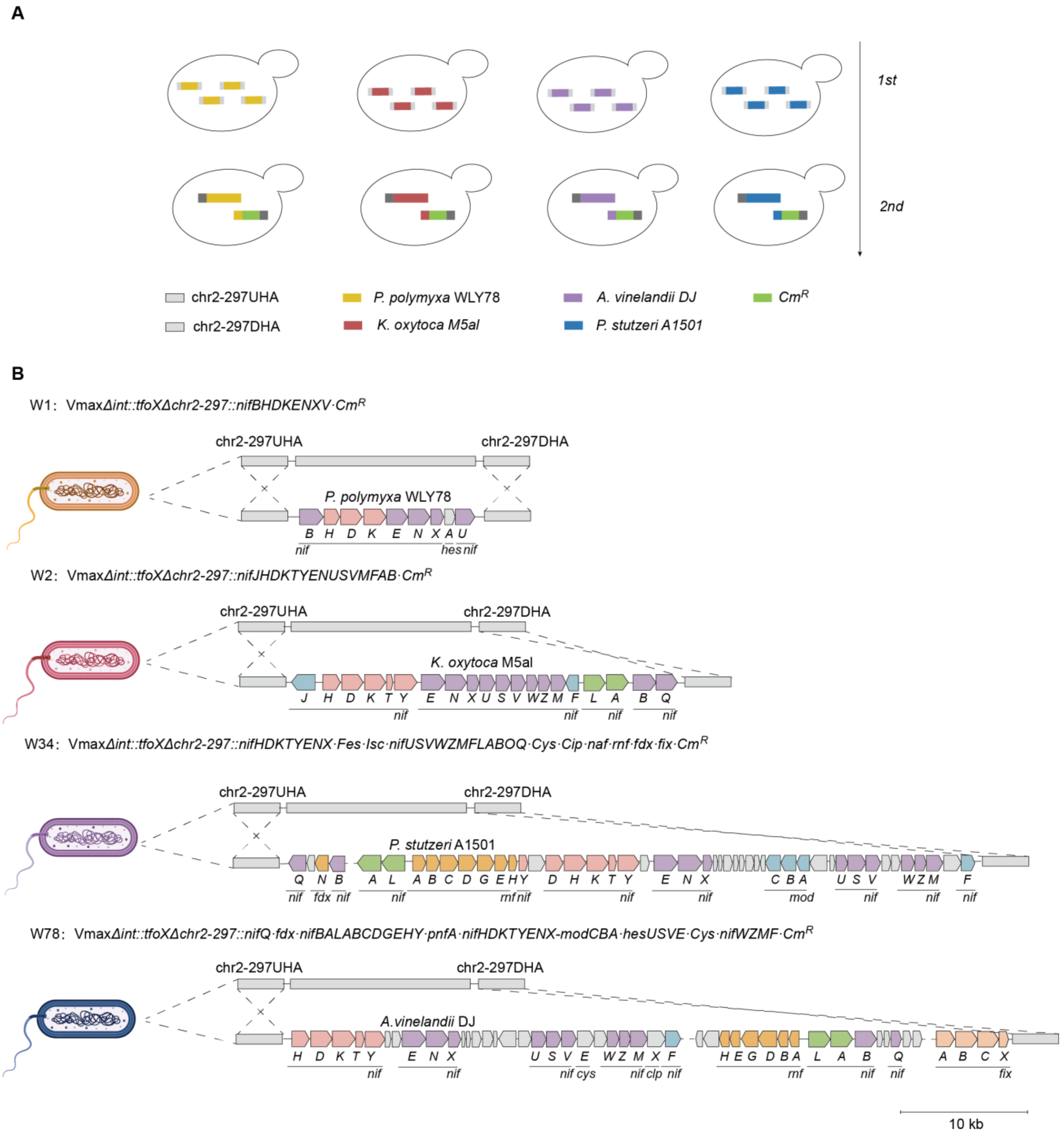


**Schematic illustration of the engineering process for ENF *V. natriegens* design.** **(A)** Assembly of *nif* gene clusters from *Paenibacillus polymyxa* WLY78, *Klebsiella oxytoca* M5al, *Azotobacter vinelandii* DJ, and *Pseudomonas stutzeri* A1501 using one-pot yeast assembly. (**B)** Integration of *nif* clusters (13-65 kb) into *V. natriegens* chromosome 2 via natural transformation, generating ENF *V. natriegens* strains W1, W2, W34, and W78.

**Fig. S2.**

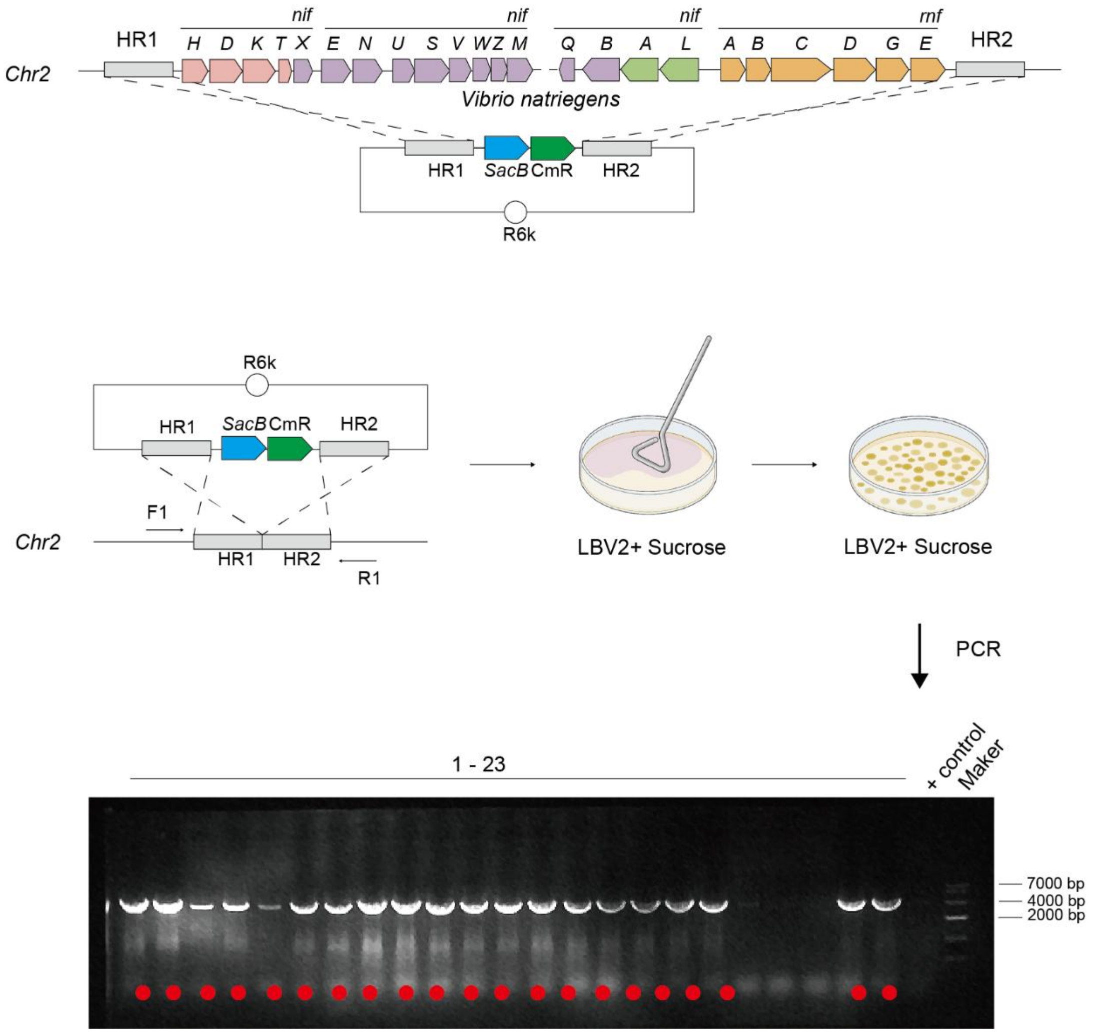


**Construction of *nif* knockout mutants in *V. natriegens*.**
Step 1: Homology arms flanking the *nif* gene cluster were cloned into a plasmid containing *SacB* and introduced via natural transformation, replacing the native *nif* cluster through homologous recombination. Step 2: The *nif* gene cluster was used to replace the antibiotic resistance marker and *SacB* via homologous recombination. Transformants were selected on LBV2 agar plates with sucrose. Step 3: PCR screening confirmed successful knockouts (red dots indicate verified clones).

**Fig. S3.**

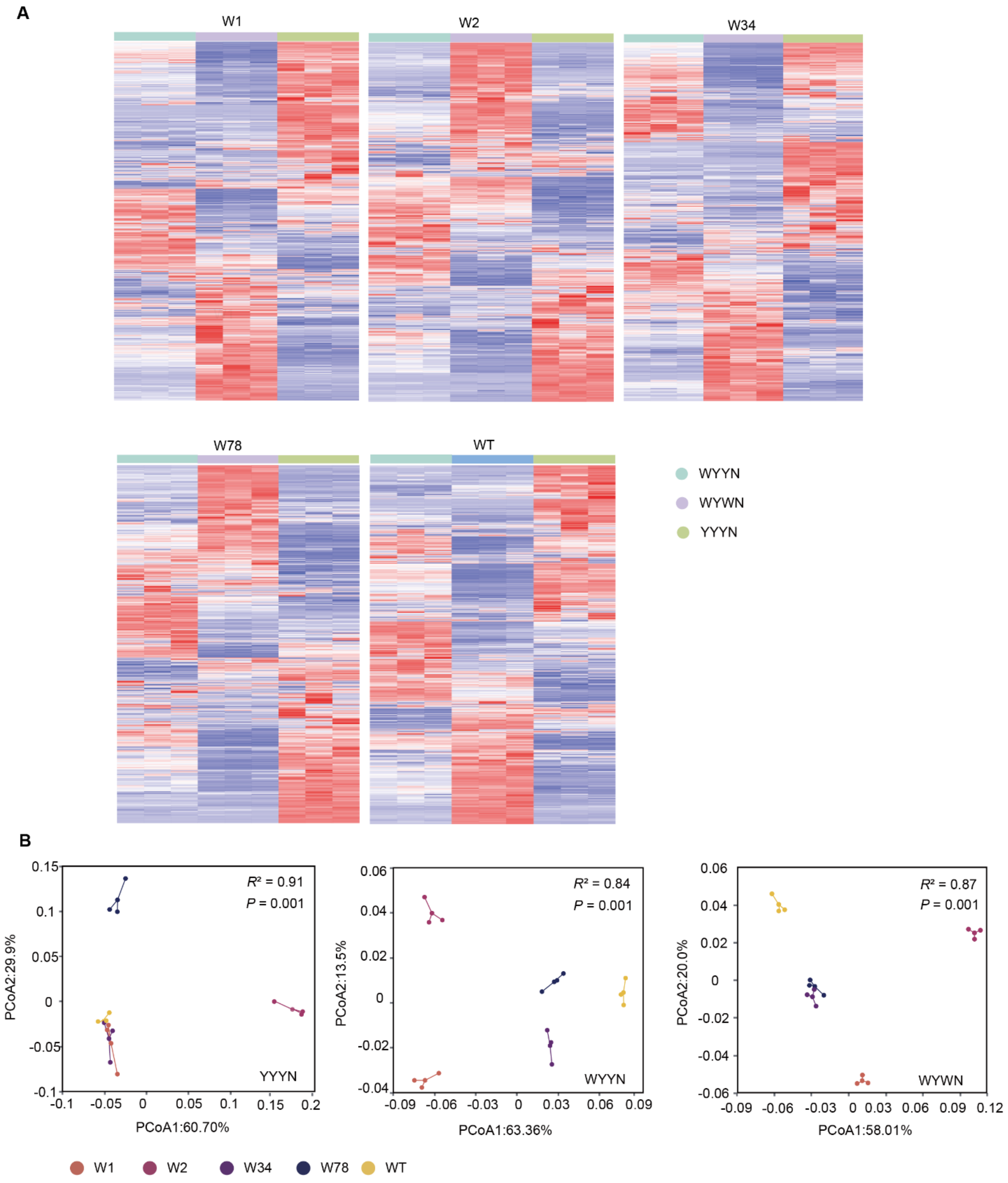


**Gene expression in ENF *V. natriegens* under different growth conditions.**
**(A)** Gene expression varies across cultivation conditions. (**B**) Principal Coordinate Analysis (PCoA) of transcriptomic profiles under different conditions: YYYN (aerobic + nitrogen), WYWN (anaerobic - nitrogen), WYYN (anaerobic + nitrogen). Statistical analysis via ADONIS: YYYN ($R^2 = 0.91, p = 0.001$); WYWN ($R^2 = 0.84, p = 0.001$); WYYN ($R^2 = 0.87, p = 0.001$).

**Fig. S4.**

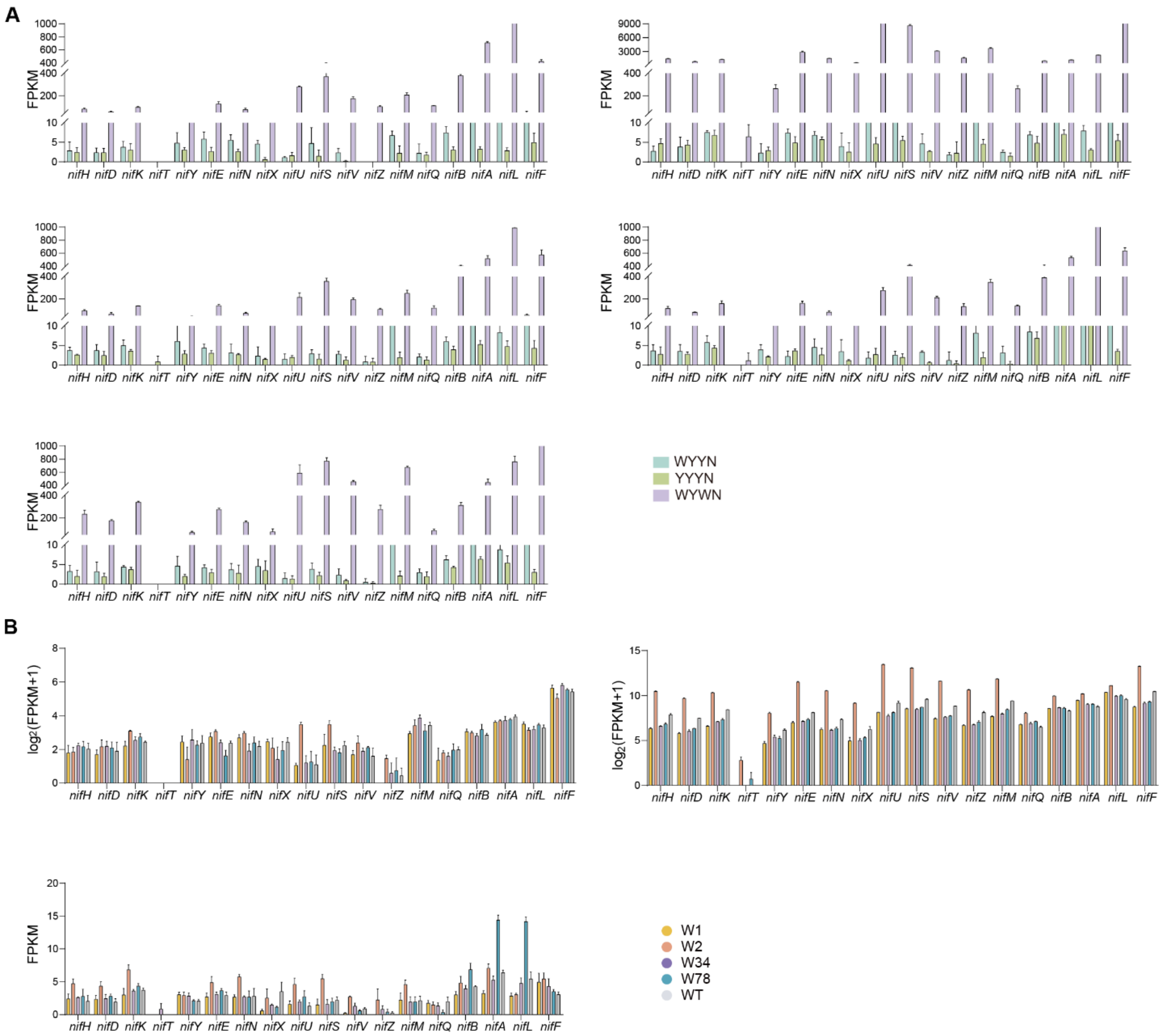


**Endogenous *nif* gene expression in ENF *V. natriegens*. (A)** Expression of endogenous *nif* genes in different ENF *V. natriegens* (W1, W2, W34, W78). **(B)** Expression of endogenous *nif* genes under different culture conditions (WYWN, YYYN, WYYN). YYYN (aerobic + nitrogen), WYWN (anaerobic - nitrogen), WYYN (anaerobic + nitrogen).

**Fig. S5.**

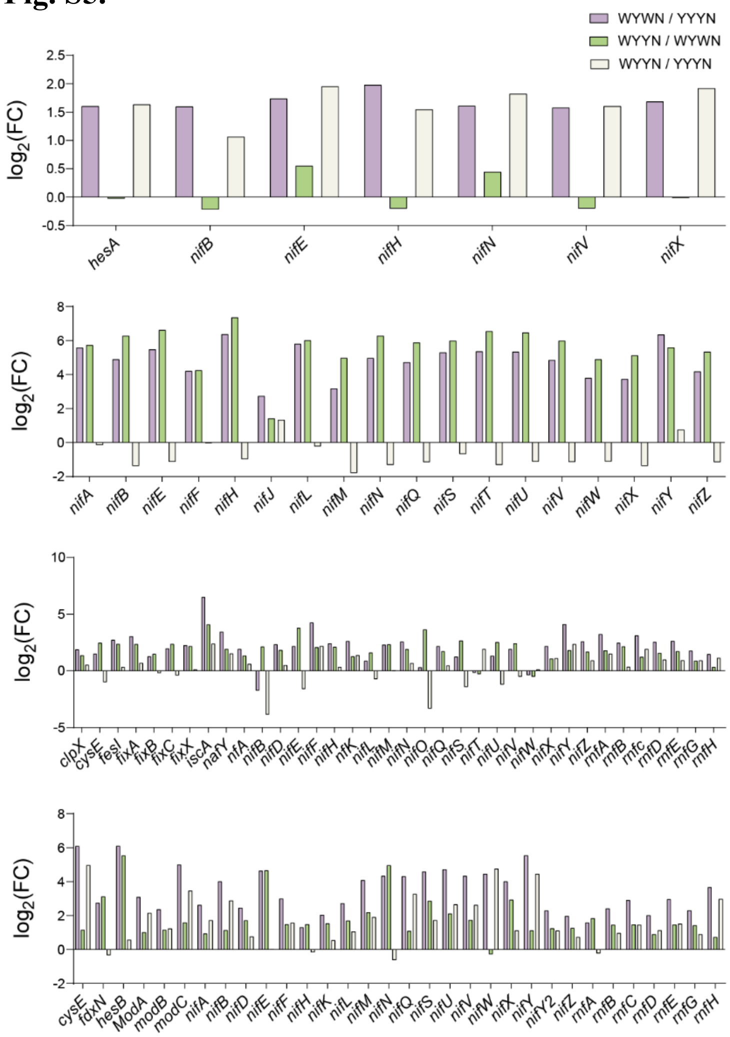


**Expression of the exogenous native *nif* gene cluster in ENF *V. natriegens* (W1, W2, W34, W78) under different culture conditions**. YYYN (aerobic + nitrogen), WYWN (anaerobic - nitrogen), WYYN (anaerobic + nitrogen). Pink: WYWN *vs*. YYYN; green: WYYN *vs*. WYWN; white: WYYN *vs*. YYYN.

**Fig. S6.**

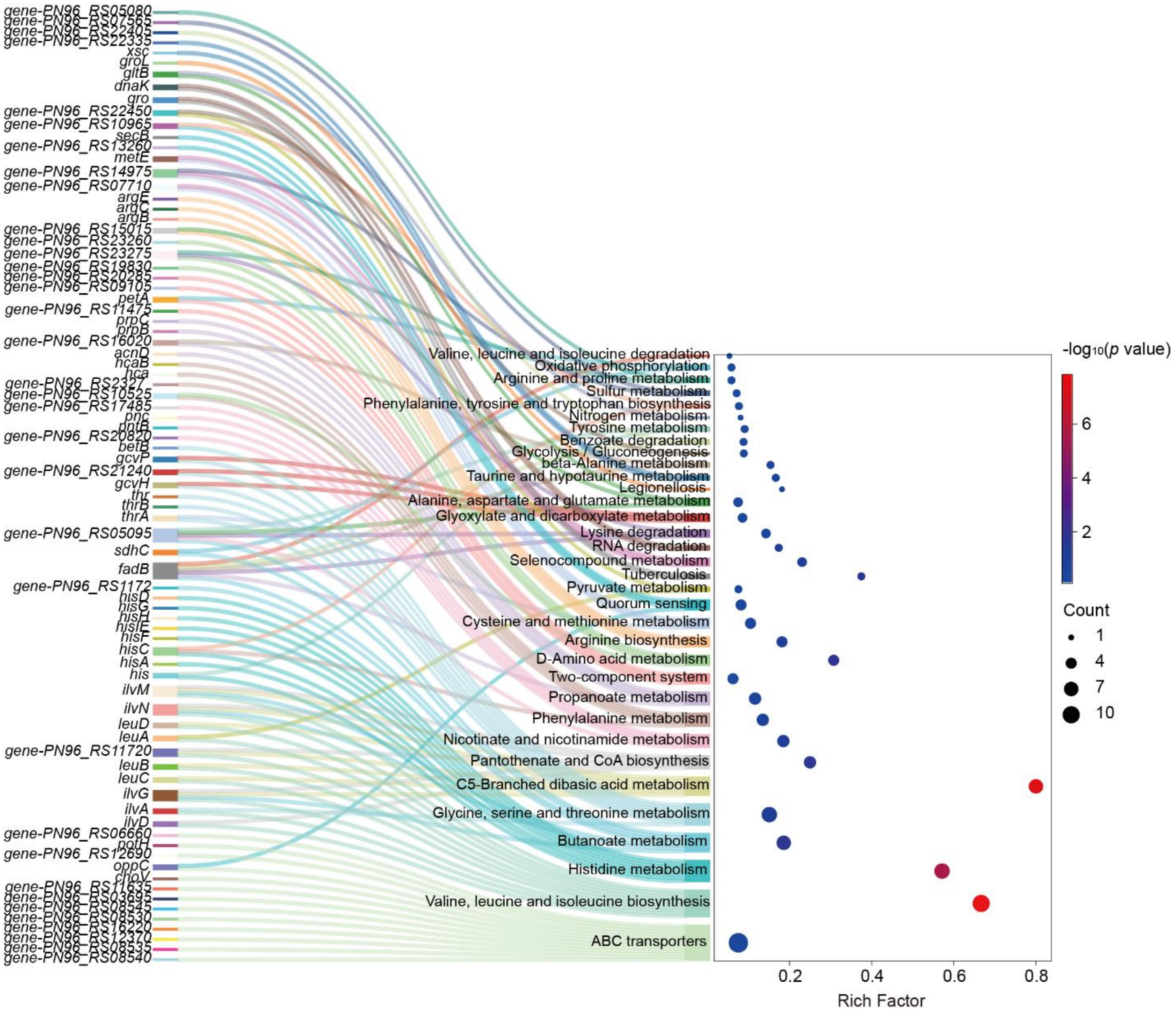


**Exogenous *nif* gene clusters enhance amino acid biosynthesis and metabolism.** A Sankey diagram shows significant upregulation of amino acid biosynthesis and metabolic pathway genes in ENF *V. natriegens* compared to the wild type *V. natriegens*.

**Fig. S7.**

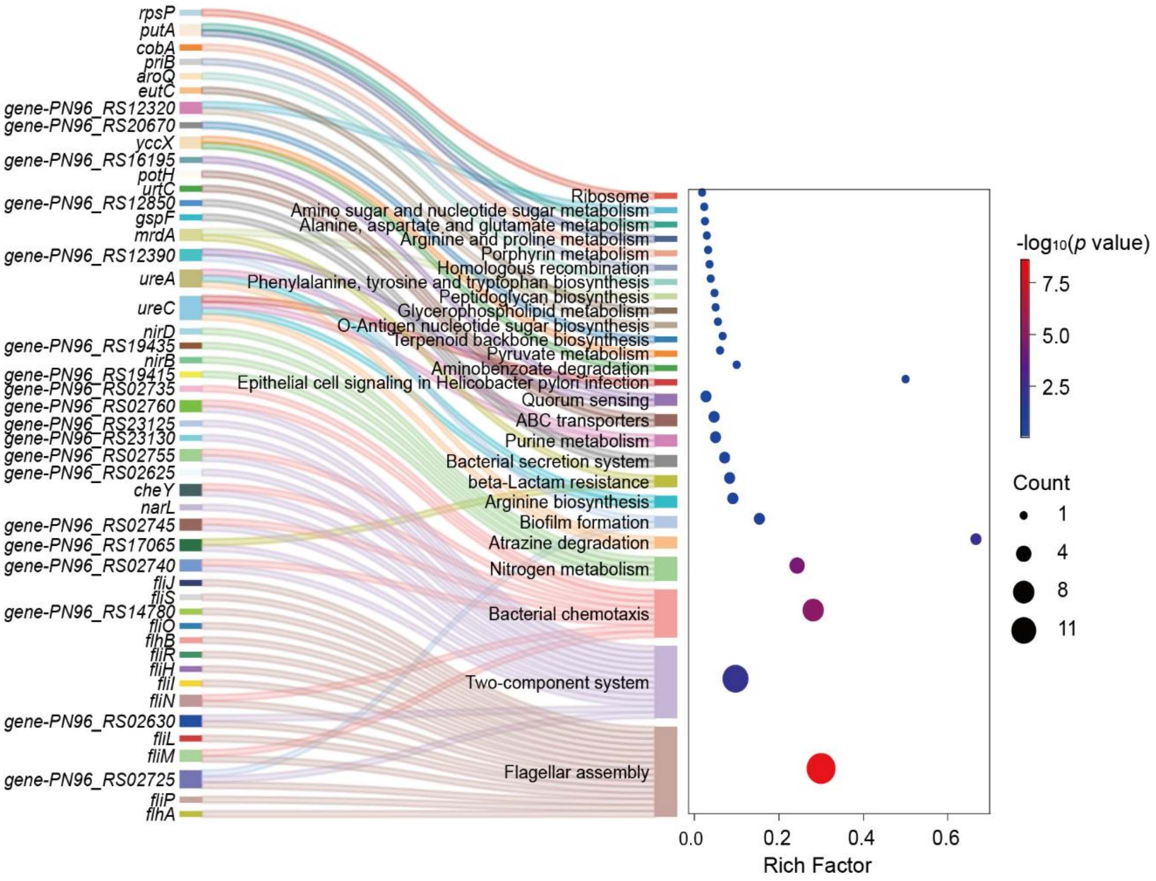


**Exogenous *nif* gene clusters modulate chemotaxis- and motility-associated gene expression.** Whole-genome analysis confirms the presence of chemotaxis- and cell motility-associated genes in *V. natriegens*. Sankey diagram shows enrichment of upregulated genes in bacterial chemotaxis and flagellar assembly pathways.

**Fig. S8.**

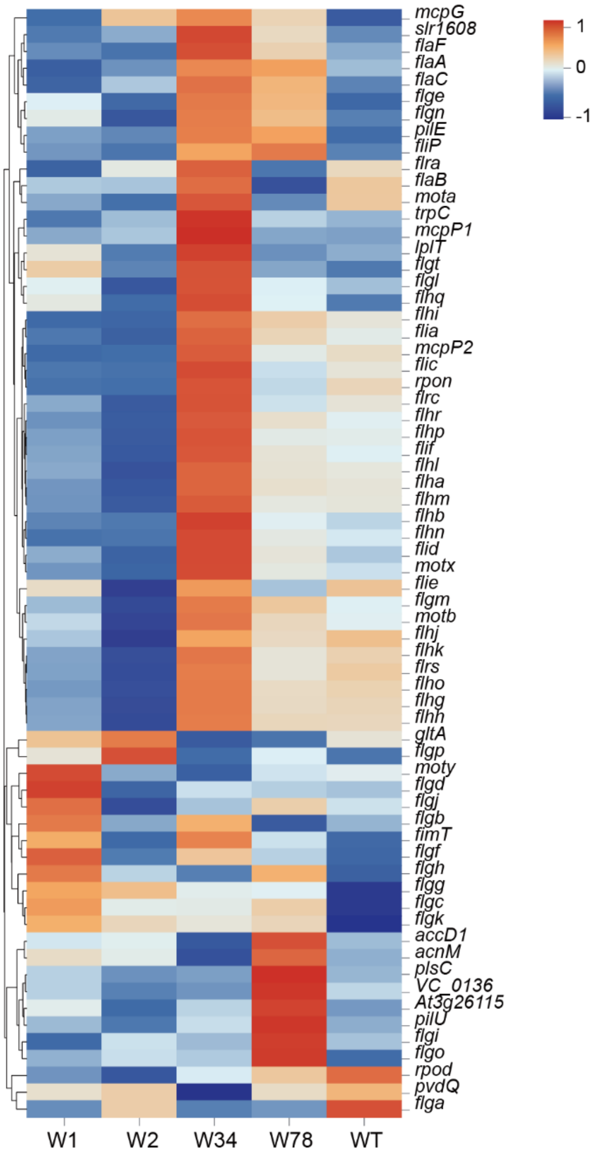


**Exogenous *nif* gene clusters enhance plant growth-promoting (PGP) genes expression in different *V. natriegens*.** The introduction of exogenous *nif* gene clusters increases the expression of plant growth-promoting (PGP) genes in different *V. natriegens*.

**Fig. S9.**

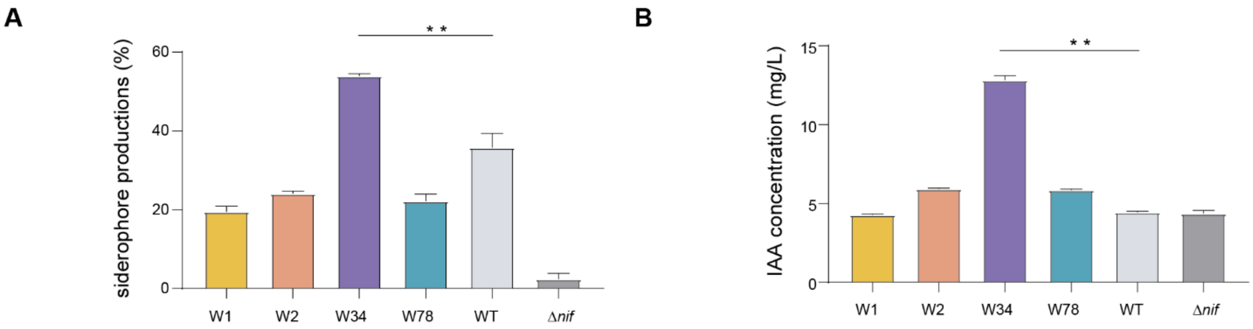


***nif*** **gene clusters enhance the secretion of plant hormones (indole-3-acetic acid, IAA) (A) and siderophores (B).**

**Fig. S10.**

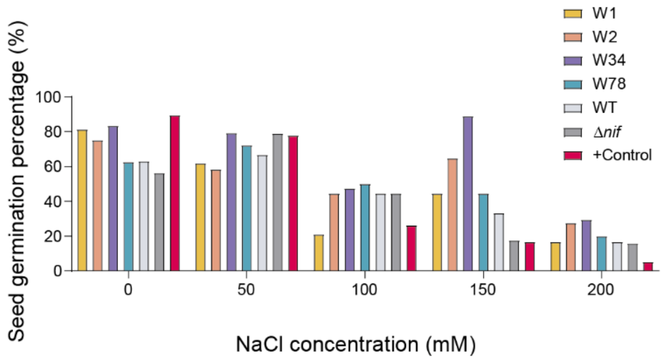


**ENF *V. natriegens* promotes soybean seed germination under different saline stress conditions.** Under high saline stress (150-200 mM NaCl), ENF *V. natriegens* W34 significantly enhances soybean germination compared to other concentrations.

**Fig. S11.**

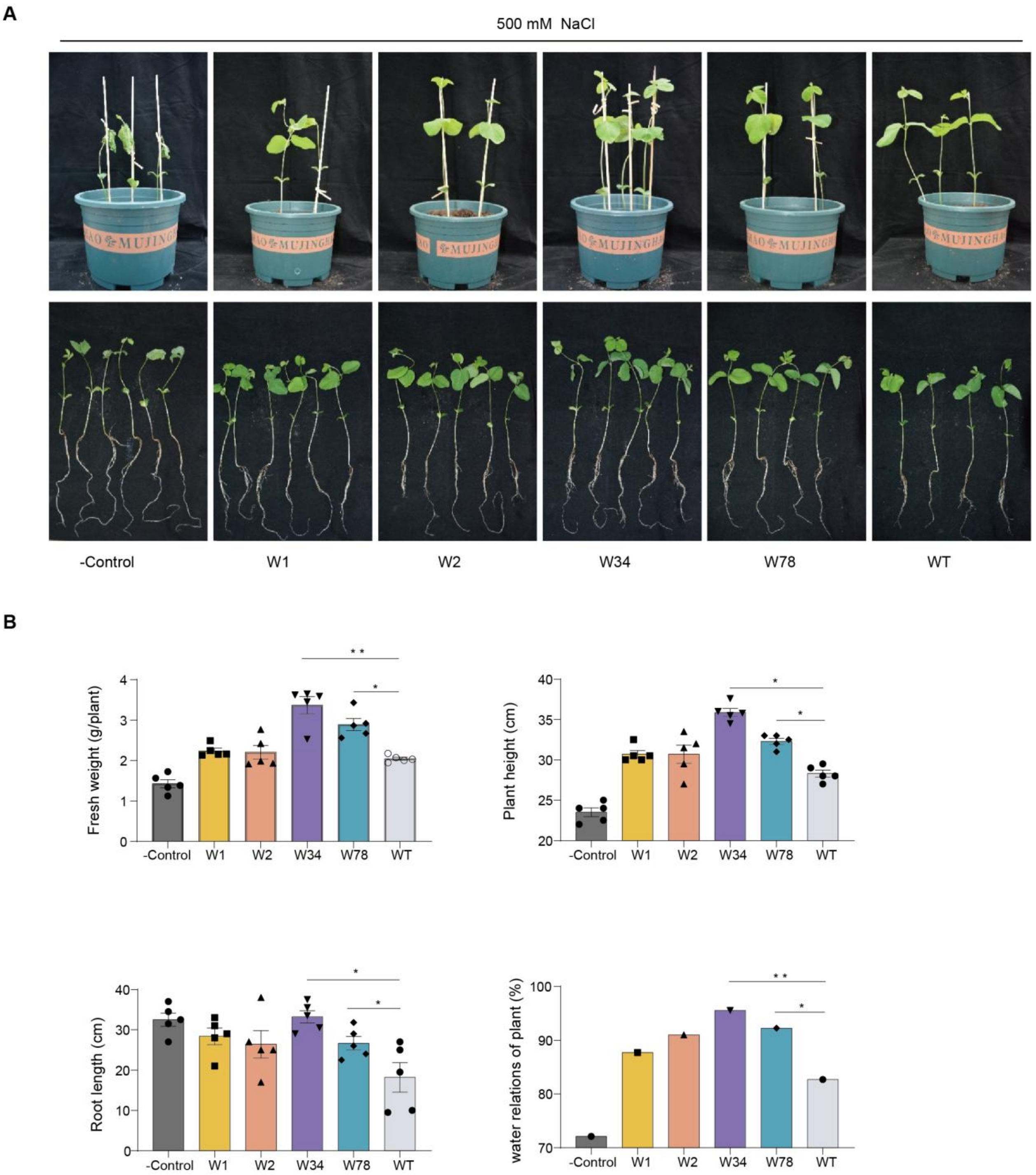


**ENF *V. natriegens* improves soybean growth under saline stress. (A)** Effect of ENF *V. natriegens* inoculation on soybean shoot phenotype under 500 mM NaCl. W34 and W78 significantly enhance seedling growth. (**B)** Plant height, fresh weight, root length, and water status vary significantly across treatments. Data are presented as mean ± SD ($n$ = 5). Error bars indicate SD. Statistical significance: *$p$ < 0.05, **$p$ < 0.01, ***$p$ < 0.001, determined by two-tailed Student's $t$-test.

**Fig. S12.**

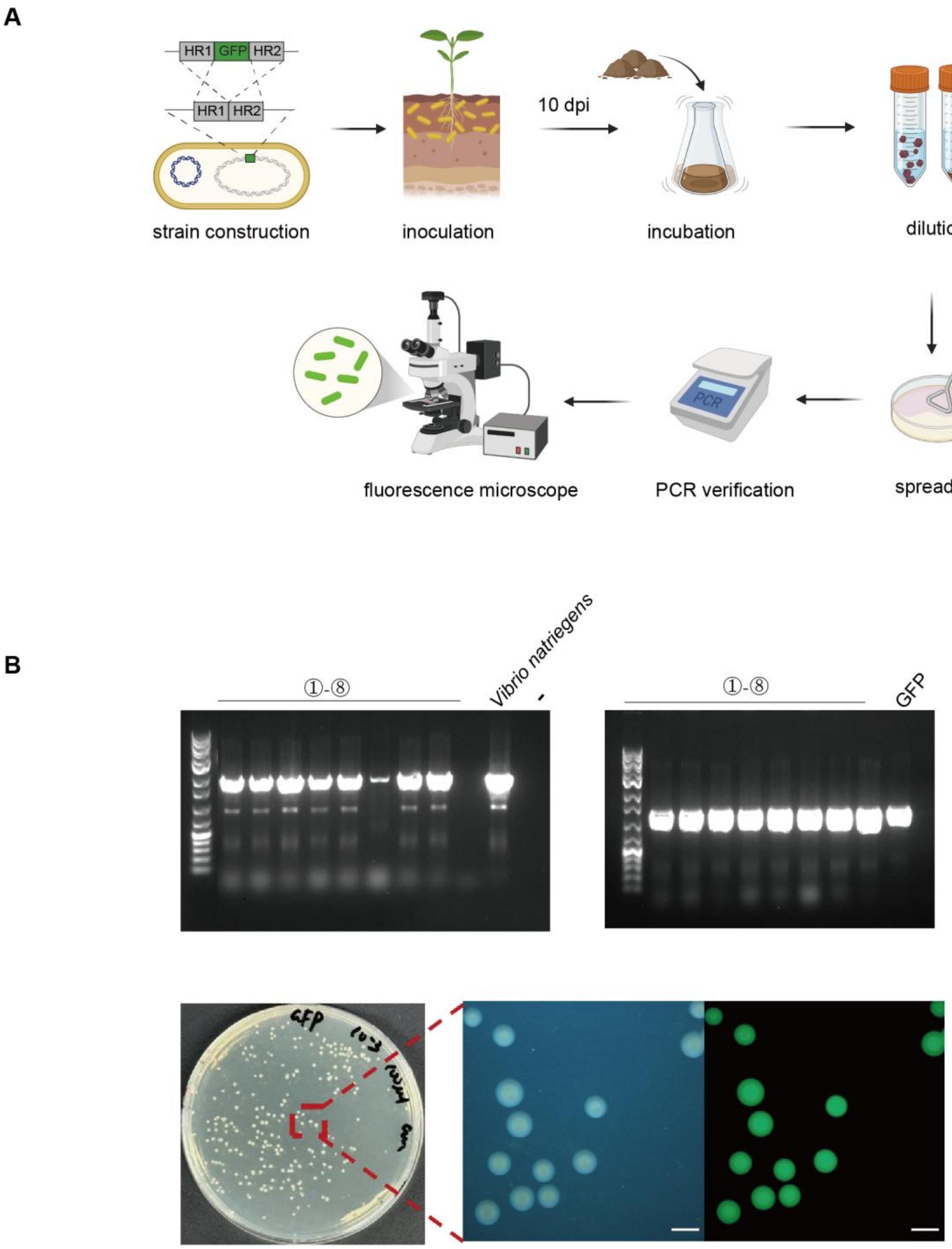


**Stable colonization of *V. natriegens* in saline soil. (A)** Schematic of the experimental workflow for observing *V. natriegens* colonization in saline soil. **Chromosomal GFP labeling**: The GFP gene was integrated into *V. natriegens* chromosome 2 via homologous recombination. **Inoculation**: GFP-tagged *V. natriegens* was introduced into the rhizosphere soil of soybean plants under saline stress. **Sampling and culture**: After 10 days, rhizosphere soil samples were mixed with M9 medium and incubated aerobically. (**B)** Serial dilutions were plated on LBV2 agar. **PCR validation**: Single colonies were PCR-validated using wild type *V. natriegens*

genomic DNA and GFP plasmid as positive controls. **Fluorescence confirmation**: GFP expression was confirmed via a fluorescence stereo microscope. Scale bar = 2000 µm.

**Fig. S13.**

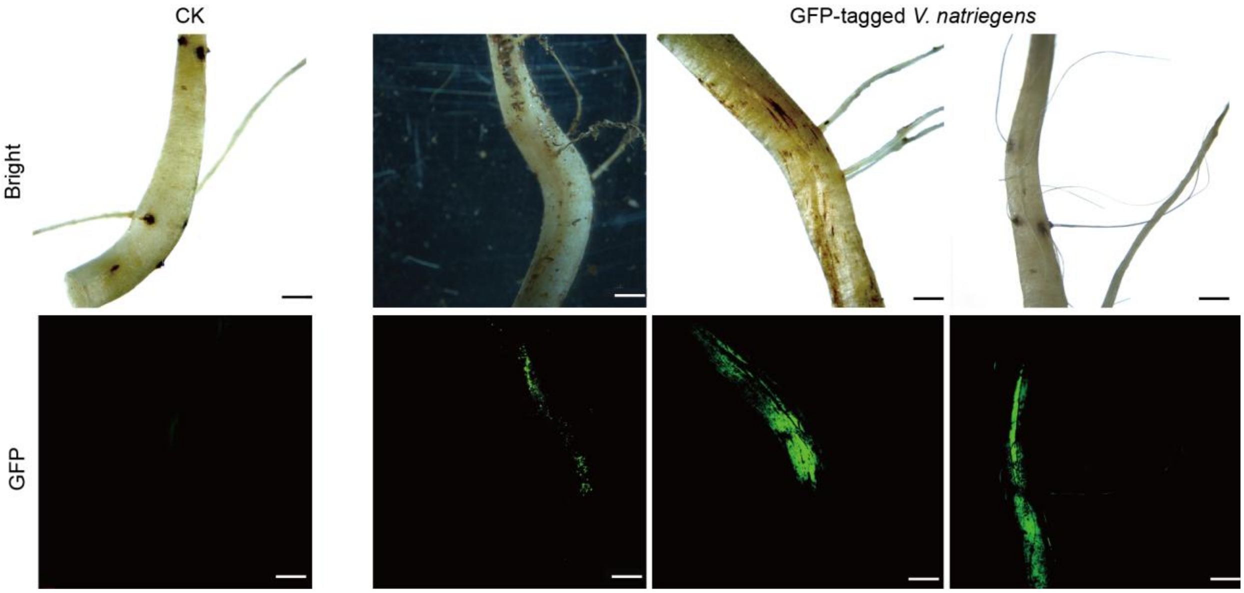


**Microscopic observation of GFP-tagged *V. natriegens* on the root surface at 10 days post inoculation (10 dpi).** Root surfaces of soybean plants inoculated with GFP-tagged *V. natriegens* were examined at 10 dpi. A total of 20 plants were analyzed under a microscope, with three biological replicates per experiment. Bright: bright-field image. Scale bar = 2000 μm.

**Fig. S14.**

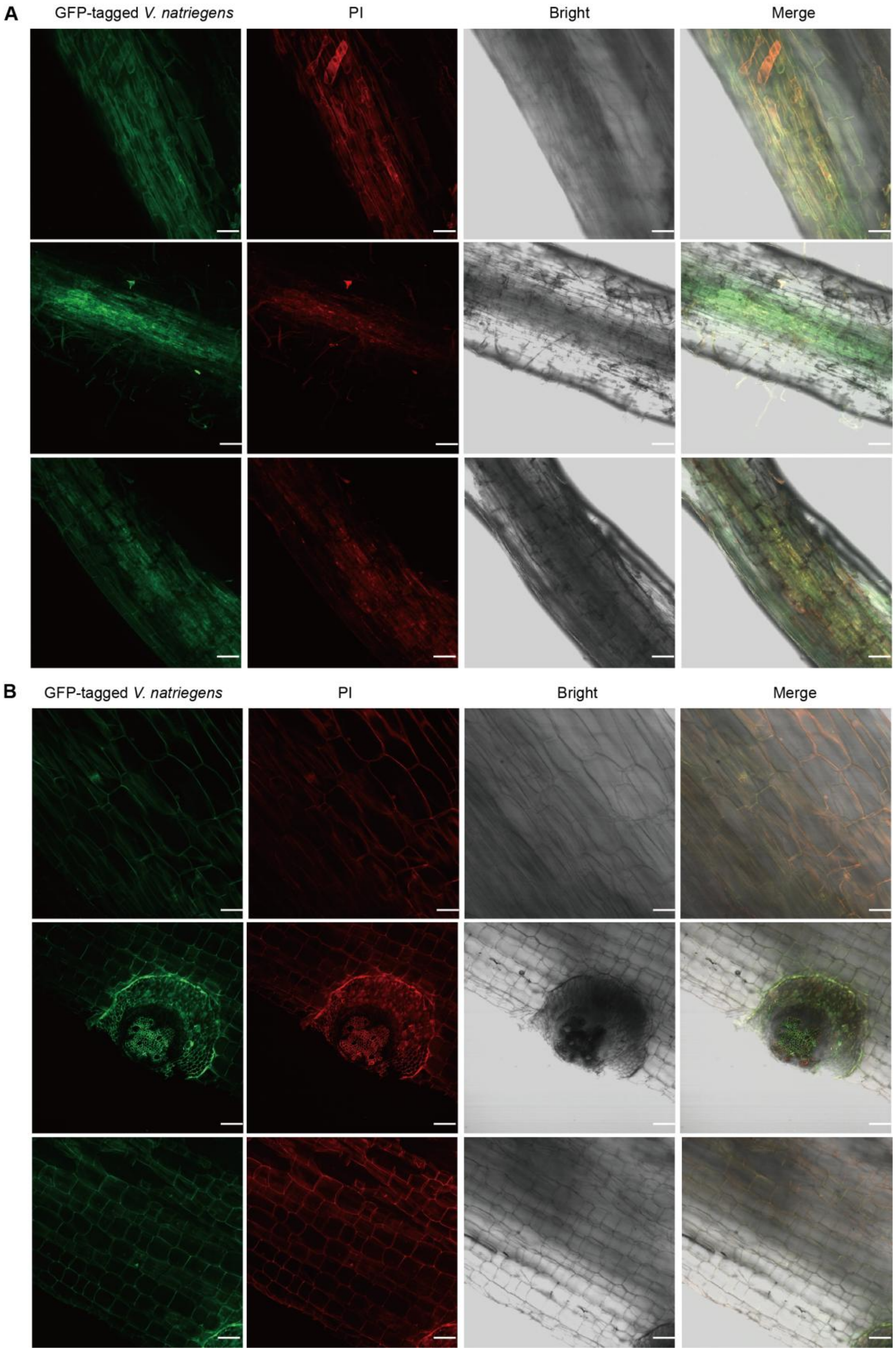

A
GFP-tagged V. natriegens
PI
Bright
Merge
B
GFP-tagged V. natriegens
PI
Bright
Merge

**Confocal laser scanning microscopy of GFP-tagged *V. natriegens* on the root surface at 10 dpi.**

(**A)** Microscopic imaging of root zones (meristem, elongation, and root hairs) inoculated with *V. natriegens*-GFP. **(B)** Confocal microscopy of longitudinal and cross-sections of roots inoculated with GFP-tagged *V. natriegens*. A total of 20 plants were analyzed with three biological replicates per experiment. PI: propidium iodide, bright: bright-field image. Scale bars = 30 μm.

**Fig. S15.**

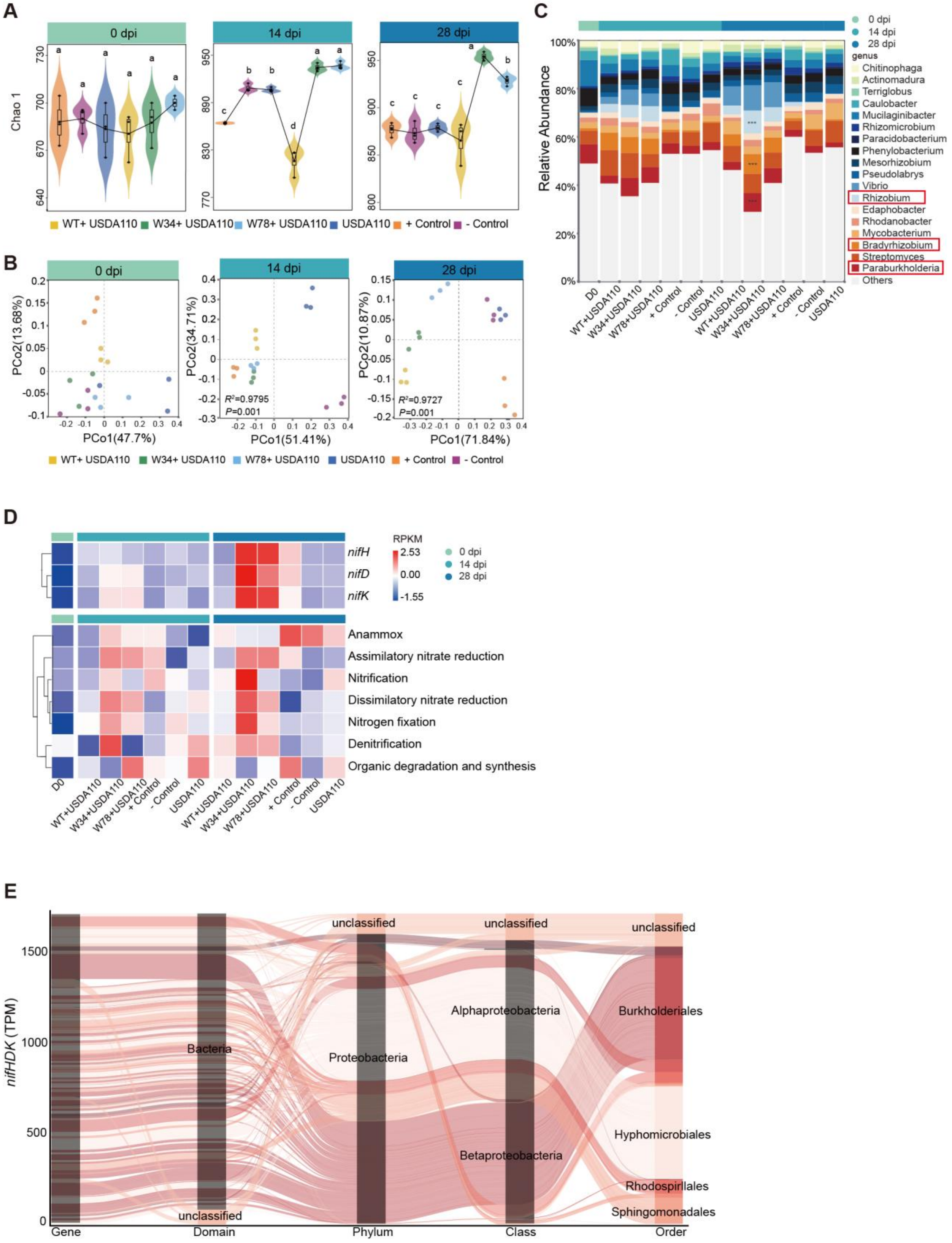
A
0 dpi
14 dpi
28 dpi
Chao 1
WT+ USDA110
W34+ USDA110
W78+ USDA110
USDA110
+ Control
- Control
B
PCo2(13.68%)
PCo1(47.7%)
PCo2(34.71%)
PCo1(51.41%)
$R^2$=0.9795
$P$=0.001
PCo2(10.87%)
PCo1(71.84%)
$R^2$=0.9727
$P$=0.001
C
Relative Abundance
genus
Chitinophaga
Actinomadura
Terriglobus
Caulobacter
Mucilaginibacter
Rhizomicrobium
Paracidobacterium
Phenylobacterium
Mesorhizobium
Pseudolabrys
Vibrio
Rhizobium
Edaphobacter
Rhodanobacter
Mycobacterium
Bradyrhizobium
Streptomyces
Paraburkholderia
Others
D0
WT+USDA110
W34+USDA110
W78+USDA110
+ Control
- Control
USDA110
D
RPKM
2.53
0.00
-1.55
nifH
nifD
nifK
Anammox
Assimilatory nitrate reduction
Nitrification
Dissimilatory nitrate reduction
Nitrogen fixation
Denitrification
Organic degradation and synthesis
E
nifHDK (TPM)
Gene
Domain
Phylum
Class
Order
Bacteria
unclassified
Proteobacteria
Alphaproteobacteria
Betaproteobacteria
Burkholderiales
Hyphomicrobiales
Rhodospirillales
Sphingomonadales

**ENF *V. natriegens* enhances nitrogen fixation activity in the rhizosphere soil.**
(**A)** Effect of different treatments on bacterial diversity (Chao 1 index) of rhizosphere soil at 0 dpi, 14 dpi, and 28 dpi under saline stress. Data are presented as mean ± SD ($n$ = 6 biologically independent samples). Different letters above the boxes indicate significant differences at $p < 0.05$ (one-way ANOVA with Tukey's HSD test).
**(B)** Principal coordinates analysis (PCoA) of 16S rRNA microbial community composition based on the Bray-Curtis dissimilarity matrix (OTU level) under different treatments at 0 dpi, 14 dpi, and 28 dpi under saline stress. Statistical analysis was performed using ADONIS (analysis of similarities) for 14 dpi (ADONIS, $R^2 = 0.9795$, $p = 0.001$) and 28 dpi (ADONIS, $R^2 = 0.9727$, $p = 0.001$).
(**C)** Relative abundance of the top 18 genera in soybean rhizosphere soil across treatments at 0 dpi, 14 dpi, and 28 dpi under saline stress. Red boxes highlight genera with significantly increased relative abundance in soybean rhizosphere soil inoculated with ENF *V. natriegens* compared to wild type *V. natriegens* at 28 dpi. Significant differences between wild type *V. natriegens* and ENF *V. natriegens* groups are indicated (*$p < 0.05$, **$p < 0.01$, ***$p < 0.001$; Student's $t$-test).
(**D)** Analyze the nitrogen cycling pathways and the expression of nitrogenase-related genes in rhizosphere soil under different treatments using functional annotation from the Ncyc database.
(**E**) Sankey plot illustrating the taxonomic distribution of *nifHDK* carriers of rhizosphere soil microbial treated with ENF *V. natriegens* W34 at the domain, phylum, class, and order levels.

**Fig. S16.**

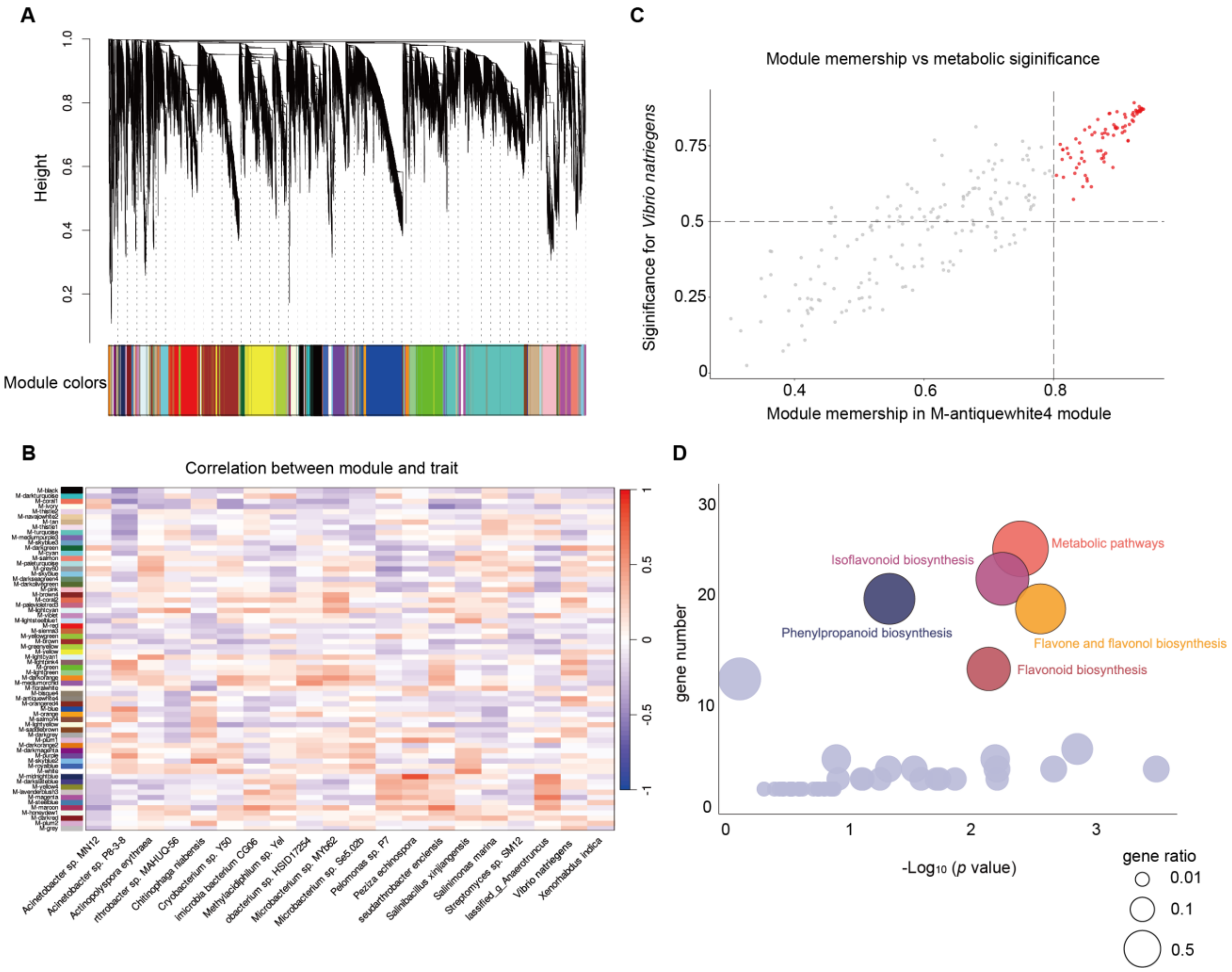


**Impact of ENF *V. natriegens* on flavonoid metabolites in soybean.**

(**A**) The relationship between species' relative abundance and differential metabolites in the gene tree was determined through weighted gene co-expression network analysis, represented by color-coded modules. Metabolites within the same module share similar expression patterns. (**B**) Microbe-module correlation heatmap: The x-axis represents different microorganisms, while the y-axis represents different metabolite modules. Each square in the matrix represents the correlation between the microorganism and the module. Red indicates a strong correlation, while blue indicates a weak correlation. Key modules were selected based on microbe-module correlation. *Pearson* correlation analysis ($R^2$ threshold = 0.8) was applied to assess associations between microbial species and differential metabolites (**C**) MM-GS analysis: The x-axis represents the module membership (MM), while the y-axis represents microbial significance (GS). Each point represents a metabolite. Red points indicate key selected metabolites. (**D**) KEGG enrichment analysis was performed on the genes in the M-antiquewhite4 module. The gene ratio in the corresponding module is represented by circle size. Light blue circles indicate pathways with lower gene ratios.

**Fig. S17.**

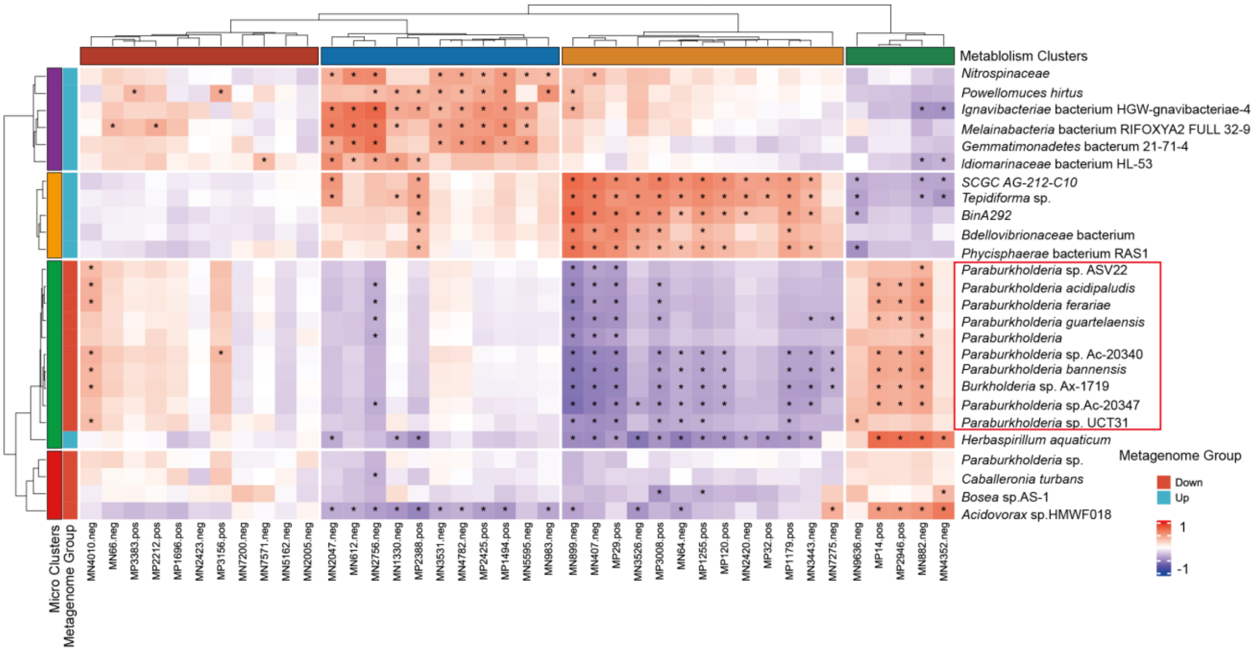


**Cluster-correlation heatmap showing associations between differential rhizosphere diazotrophs and flavonoid metabolites in soybean.** Hierarchical clustering groups co-varying microbial taxa and metabolites into color-coded clusters (labeled boxes), revealing modular interactions between nitrogen-fixing microbiota and plant secondary metabolism. Microbial taxa (horizontal axis) and metabolites (vertical axis) are organized based on similarity in correlation patterns. Color intensity represents *Spearman* correlation coefficients (red: positive; blue: negative), with asterisks indicating significant correlations ($p < 0.05$).

**Fig. S18.**

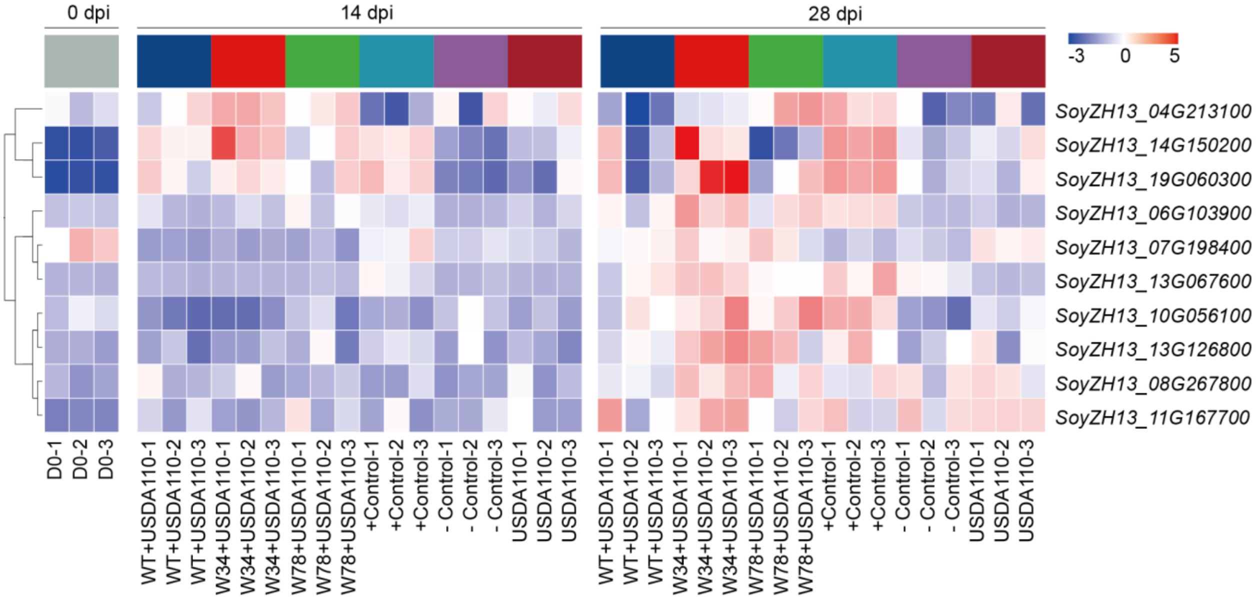


**Expression of nitrogen metabolism-related genes in soybean under different treatments.**

**Fig. S19.**

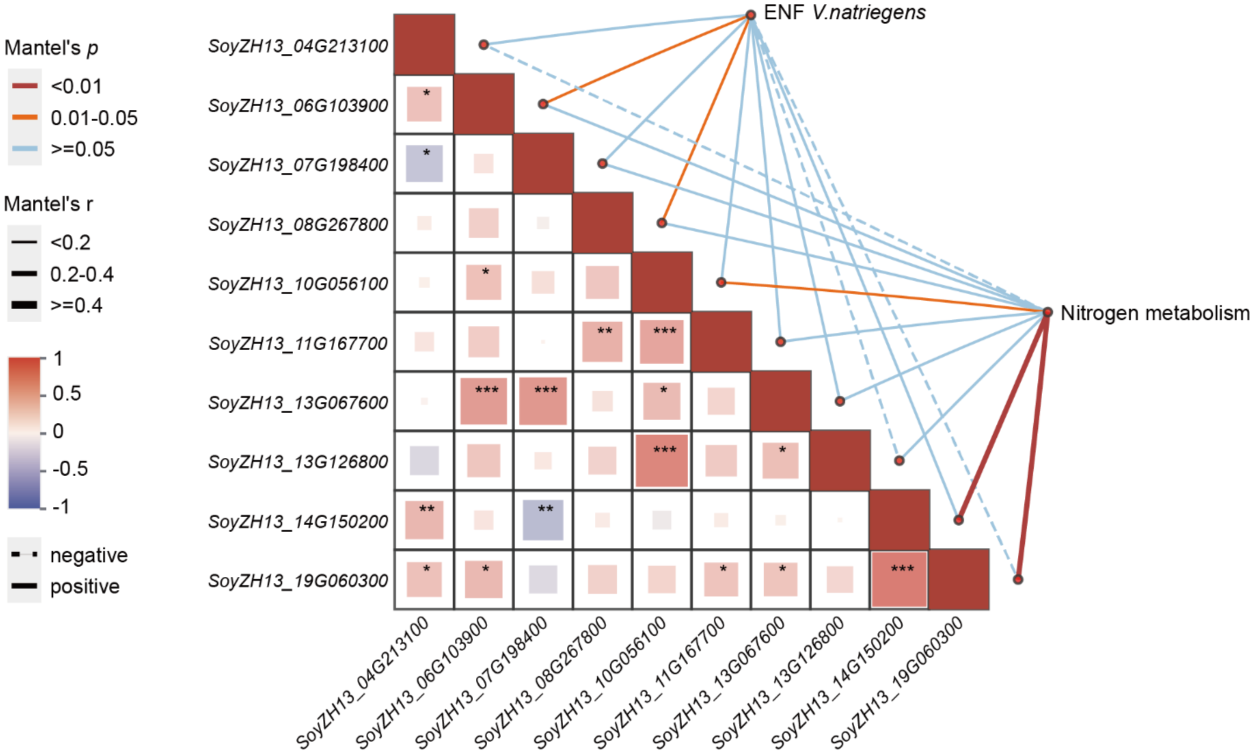


**Relationships between ENF *V. natriegens abundance*, nitrogen metabolism, and nitrogen-related genes using *Mantel* tests.** *Mantel* tests were performed to analyze correlations based on *Pearson*'s coefficients. Line width represents the magnitude of the absolute value of *Mantel*'s r, and line color indicates significance levels (*$p < 0.05$, **$p < 0.01$, ***$p < 0.001$).

**Fig. S20.**

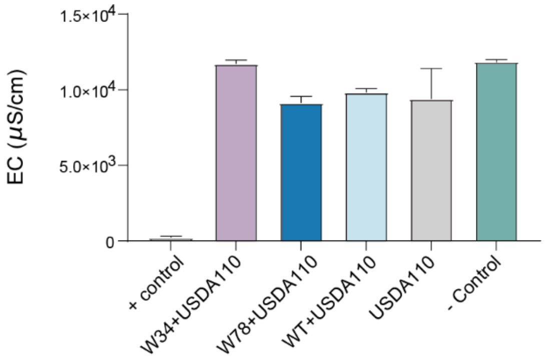


**The electrical conductivity of rhizosphere soil under different treatments.**

**Fig. S21.**

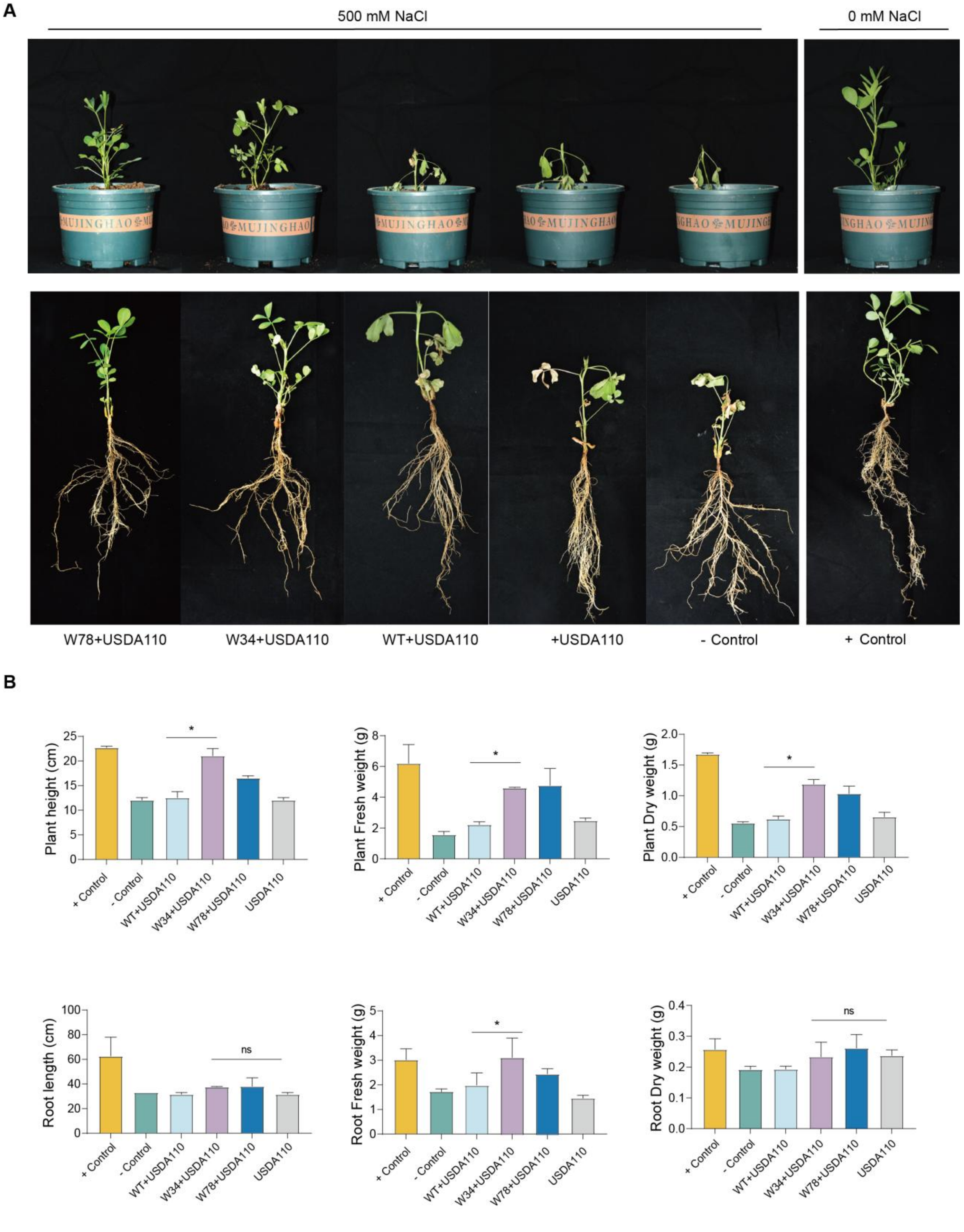
A
500 mM NaCl
0 mM NaCl
W78+USDA110
W34+USDA110
WT+USDA110
+USDA110
- Control
+ Control
B
Plant height (cm)
Plant Fresh weight (g)
Plant Dry weight (g)
Root length (cm)
Root Fresh weight (g)
Root Dry weight (g)
+ Control
- Control
WT+USDA110
W34+USDA110
W78+USDA110
USDA110
*
ns

**Impact of ENF *V. natriegens* on peanut growth under saline stress. (a)** Phenotypic effects of *V. natriegens* inoculation under 500 mM saline stress. The addition of W34 and W78 significantly improved peanut growth under saline stress. (**b)** Plant growth parameters under 500 mM saline stress. Significant differences were observed in height, fresh weight, dry weight, root length, root fresh weight, and root dry weight. Data are presented as the mean ± SD ($n$ = 10). Error bars indicate the standard deviations of ten replicates (*$p$ < 0.05, **$p$ < 0.01, ***$p$ < 0.001; Student's $t$-test).

**Fig. S22.**

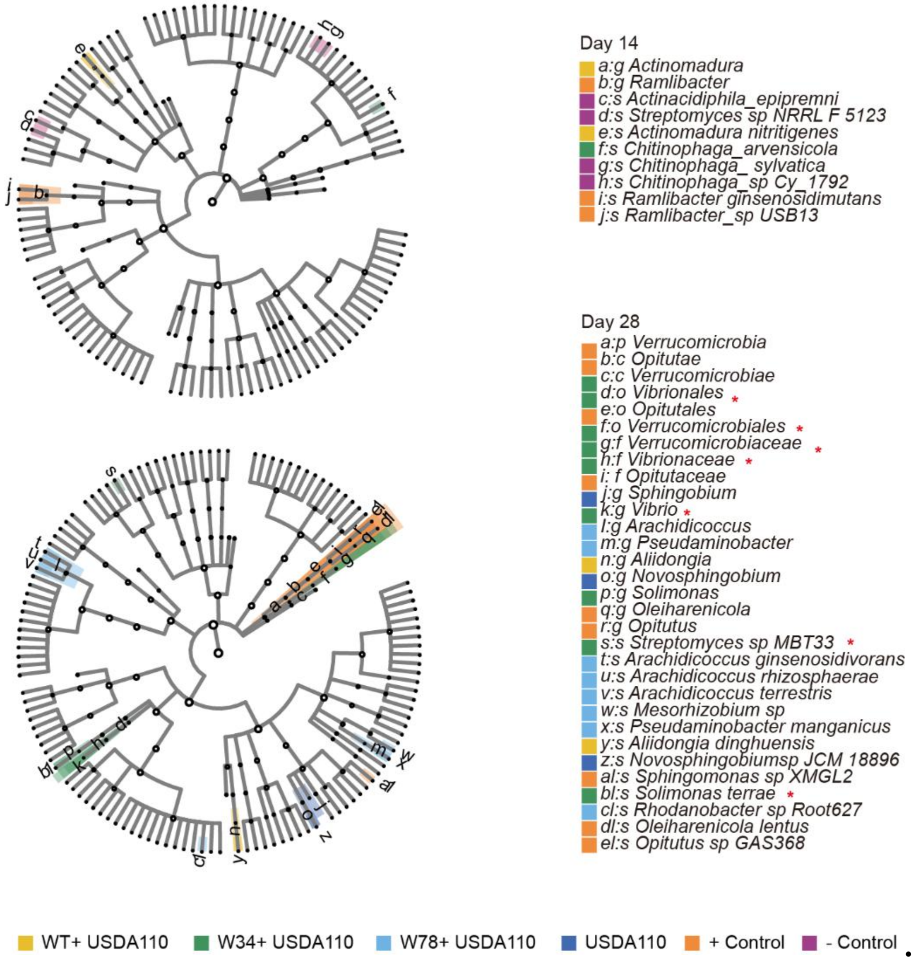


***LefSe* analysis of rhizosphere soil microbiota under different strain treatments at different soybean developmental stages (0 dpi, 14 dpi, 28 dpi)**. *LefSe* identified significantly different bacterial taxa based on biomarkers with LDA scores > 4. The radial diagram represents classification levels from phylum to species. Each small circle represents a taxonomic classification, with its diameter corresponding to relative abundance. Different colors indicate different treatment groups, and nodes of different colors represent key microbial taxa within each group. Red asterisks denote microbial taxa that show significant differences in rhizosphere soil inoculated with ENF *V. natriegens* W34 compared to other treatment groups.

**Table S1. Primers used for *nif* gene cluster cloning.**

| *nif* cluster | Forward primer | Reverse primer | GenBank accession No. |
|---|---|---|---|
| *Paenibacillus polymyxa* WLY78 | GAATTGAGGATAAATGTCAGGGATTTCATG | ACAGGTTCCGCAGTTCACAAGC | ALJV01.1 contig00089 |
| | CCAAGCATTTTGAGATCGCGGATG | GCTGATTGTGATCGACAATATTCGG | |
| | CGGAGGTGCCGGTATGAGCGA | GAAAGCCTACACGAAGCAAAGG | |
| | GAAGTTTGCAGCGAAAGAGGCG | TTAATCAGCAAATTCCGTCTCCTCC | |
| | GGGATGATGCAGAATACATCCCG | ATCCACAAATCAACACCCTGCG | |
| | GGTGACCTGGATGATGCAGAGGAGAG | AAAGCGTTCCAGTCACGGTCAC | |
| *Klebsiella oxytoca* M5al | CTGCTGGATACGCTGCTTAAGGTC | GTGACGCTCGCGTATCAGGTTTG | CP020657.1 |
| | TACGCTGTTTGAGCTGGCAAACCT | GGTTGCGGAAGCTTTTGGTCAC | |
| *Azotobacter vinelandii* DJ | GGGCAGCCAGTGGAAAAAG | TTCGTCAGTGGAGTGGCTTGC | NC_012560 |
| *Pseudomonas stutzeri* A1501 | GTTAGGTTGGCCTGAATTCGGTG | GAACTCATGGCGCAACATGCG | NC_009434 |
| | GATGACCATGAGAAAGATCGACGT | CATGAGCGAGGATGGCACC | |
| | GAATGTTCAGCGTCTGCACTCG | GTTGACCAGCACGGTCTTGAT | |
| | CATCCTCTCGATCGGGCCG | CTTTAACGGCATGTTCCGGGTC | |
| | CTGCACGAGTGGCATGTCA | CGATCACGGATGTCGGCAG | |
| | GAAGGCCGAGGAATC | GTAGAAGCTGCCCTC | |

| | | |
|---|---|---|
| | GGC | GAACCAG |
| | CGCCGAGGTGAACATGATGG | GGGCTGAAGGCGAAGCC |
| | GAGCTCTACGACTGGCTGC | GACCAGCGCACGCAATCA |
| | CAACTGCAGTTCGTCGATGCTG | CATTAGCACCGATGCTGATCGG |
| | GACGACGTCACCCTCTATCACG | GGCGAATCTCCTTCCTCGG |

**Table S4. PGPB with their saline tolerance values.**

| PGPB | Crops | Salt level | Ref. |
|---|---|---|---|
| *Pseudomonas putida* 108, *Pseudomonas fluorescens* 153, *Pseudomonas chlororaphis* TSAU13, *Pseudomonas extremorientalis* TSAU20 | Wheat, Bean, Cotton | 12 dS $m^{-1}$ | 5 |
| *Azospirillum brasilense* NH | Wheat | 200 mM NaCl | 6 |
| *Achromobacter piechaudii* ARV8 | Tomato | 172 mM NaCl | 7 |
| *Streptomyces venezuelae* ATC | Thai rice | 150 mM NaCl | 8 |
| *Bradirhizobium Japanicum* | Soybean | 61 to 75mMNaCl | 9 |
| *Pseudomonas simiae* | Soybean | 100 mM NaCl | 10 |
| *Pseudomonas pusedoalcaligenes* | Rice | 2.5 g NaCl $kg^{-1}$soil | 11 |
| *Bacillus amyloliquefaciens* NBRISN13 | Rice | 200 mM NaCl | 12 |
| *Brevibacterium* l*inens* RS16 | Rice | 100 mM NaCl | 13 |
| *Pseudomonas frederiksbergensis* OS261 | Red pepper | 150 mM of NaCl | 14 |
| *Enterobacter* sp. UPMR18 | Okra | 75 mM NaCl | 15 |
| *Co-inoculation of Rhizobium phaseoli & P. fluorescens* | Mungbean | 4.1–6.7 dS $m^{-1}$ | 16 |
| *Bacillus megaterium* | Maize | 150 mM NaCl | 17 |
| *Pantoea agglomerans* | Maize | 0.2 mol l−1NaCl | 18 |
| *Pseudomonas putida* AKMP7 | Maize | 100 mM NaCl | 19 |
| *Rhizobium* (strain Thal-8) and *phosphate-solubilizing bacteria (Pseudomonas* sp. 54RB*)* applied alone and in combination | Maize | 100 mM NaCl | 20 |
| *Bacillus amyloliquefaciens* SQR9 | Maize | 100 mM NaCl | 21 |
| *Azospirillum brasilense* Sp245 Sp | Lettuce | 40 mM NaCl | 22 |
| *P. fluorescens* | Groundnut | 120 mM NaCl | 23 |
| *Rhizobium leguminosarum vicia* strain GRA 19, 3841, *Bradyrhizobium* sp. | Faba bean | 100–120 mM NaCl | 24 |
| *Pseudomonas putida* UW4 | Canola | 100 mM NaCl | 25 |
| *Sinorhizobium mellilote* R29, R103*, Rhizobium* legominozarum *bv. Phaseoli strains* R281, R307 | Canola | 50 mM NaCl | 26 |
| *Bacillus subtilis* GB03 | Arabidopsis | 100 mM NaCl | 27 |
| *Streptomyces* sp. *strain* PGPA39 | Tomato | 180 mM NaCl | 28 |

**Table S5. Strains used in this study**

| name | strain | source | description |
|---|---|---|---|
| Vmax | *V. natriegen*s | This study | |
| USDA110 | *Bradyrhizobium diazoefficiens* | Tian lab | |
| A1501 | *P. stutzeri* | This study | |
| BY4742 | *Saccharomyces cerevisiae* | Dai lab | |
| EPI300 | *E. coli* | Dai lab | |
| pRS415 | *E. coli* | Dai lab | |
| pJS356 | *E. coli* | Dai lab | |
| WWE001 | *E. coli* | This study | *P. polymyxa* WLY78 |
| WWE002 | *E. coli* | This study | *K. oxytoca* M5al |
| WWE003 | *E. coli* | This study | *A. vinelandii* DJ-1 |
| WWE004 | *E. coli* | This study | *A. vinelandii* DJ-2 |
| WWE007 | *E. coli* | This study | *P. stutzeri* A1501-1 |
| WWE008 | *E. coli* | This study | *P. stutzeri* A1501-2 |
| WWE001UD | *E. coli* | This study | *P. polymyxa* WLY78-UD |
| WWE002UD | *E. coli* | This study | *K. oxytoca* M5al-UD |
| WWE003UD | *E. coli* | This study | *A. vinelandii* DJ-1-UD |
| WWE034 | *E. coli* | This study | *A. vinelandii* DJ |
| WWE034UD | *E. coli* | This study | *A. vinelandii* DJ-UD |
| WWE078 | *E. coli* | This study | *P. stutzeri* A1501 |
| WWE078UD | *E. coli* | This study | *P. stutzeri* A1501-UD |
| WWY001 | *Saccharomyces cerevisiae* | This study | *P. polymyxa* WLY78 |
| WWY002 | *Saccharomyces cerevisiae* | This study | *K. oxytoca* M5al |
| WWY003 | *Saccharomyces cerevisiae* | This study | *A. vinelandii* DJ-1 |
| WWY004 | *Saccharomyces cerevisiae* | This study | *A. vinelandii* DJ-2 |
| WWY007 | *Saccharomyces cerevisiae* | This study | *P. stutzeri* A1501-1 |
| WWY008 | *Saccharomyces cerevisiae* | This study | *P. stutzeri* A1501-2 |
| WWY001UD | *Saccharomyces cerevisiae* | This study | *P. polymyxa* WLY78-UD |
| WWY002UD | *Saccharomyces cerevisiae* | This study | *K. oxytoca* M5al-UD |
| WWY003UD | *Saccharomyces cerevisiae* | This study | *A. vinelandii* DJ-1-UD |
| WWY034 | *Saccharomyces cerevisiae* | This study | *A. vinelandii* DJ |
| WWY034UD | *Saccharomyces cerevisiae* | This study | *A. vinelandii* DJ-UD |
| WWY078 | *Saccharomyces cerevisiae* | This study | *P. stutzeri* A1501 |
| WWY078UD | *Saccharomyces cerevisiae* | This study | *P. stutzeri* A1501-UD |
| W1 | *V. natriegen*s | This study | *P. polymyxa* WLY78 |
| W2 | *V. natriegen*s | This study | *K. oxytoca* M5al |
| W34 | *V. natriegen*s | This study | *A. vinelandii* DJ |
| W78 | *V. natriegen*s | This study | *P. stutzeri* A1501 |
| Vmax*Δnif* | *V. natriegen*s | This study | |
| Vmax::GFP | *V. natriegen*s | This study | |

**Table S6. Primers used for real-time PCR in this study[29]**

| *nif* cluster | primer |
|---|---|
| SoyZH13_13G126800-F | TGAGAGTGGTGCCAAAGTTAG |
| SoyZH13_13G126800-R | CATGGCCTCTGTTCTTGATAGT |
| SoyZH13_03G215600-F | CTTCTCGTGGGACTAGGAATTG |
| SoyZH13_03G215600-R | CCATCCCTTCTTCTATGTCTTTCT |
| Actin 11 -F | CCATCCCTTCTTCTATGTCTTTCT |
| Actin 11 -R | CGTTTCATGAATTCCAGTAGC |

**Table S7. Composition of nutrient solution for soybean growth**

| Element | Final Molarity (µM) | Chemical Form | Molecular Weight | Mass per Liter (g) | Stock Solution Molarity |
|---|---|---|---|---|---|
| Ca | 1000 | $CaCl_2$-$2H_2O$ | 147.03 | 294.1 | 2 |
| P | 500 | $KH_2PO_4$ | 136.09 | 136.1 | 1 |
| Fe | 10 | Fe-Citrate | 335.04 | 6.7 | 0.02 |
| Mg | 250 | $MgSO_4$-$7H_2O$ | 246.5 | 123.3 | 0.5 |
| K | 1500 | $K_2SO_4$ | 174.06 | 87 | 0.5 |
| Mn | 1 | $MnSO_4$-$H_2O$ | 169.02 | 0.338 | 0.002 |
| B | 2 | $H_3BO_4$ | 61.84 | 0.247 | 0.004 |
| Zn | 0.5 | $ZnSO_4$-$7H_2O$ | 287.56 | 0.288 | 0.001 |
| Cu | 0.2 | $CuSO_4$-$5H_2O$ | 249.69 | 0.1 | 0.004 |
| Co | 0.1 | $CoSO_4$-$7H_2O$ | 281.12 | 0.056 | 0.0002 |
| Mo | 0.1 | $Na_2MoO_4$-$2H_2O$ | 241.98 | 0.048 | 0.0002 |
